# Influence of expected reward on perceptual decision making

**DOI:** 10.1101/506790

**Authors:** Mohsen Rakhshan, Vivian Lee, Emily Chu, Lauren Harris, Lillian Laiks, Peyman Khorsand, Alireza Soltani

## Abstract

Perceptual decision making is influenced by reward expected from alternative options or actions, but the underlying neural mechanisms are currently unknown. More specifically, it is debated whether reward effects are mediated through changes in sensory processing and/or later stages of decision making. To address this question, we conducted two experiments in which human subjects made saccades to what they perceived to be the first or second of two visually identical but asynchronously presented targets, while we manipulated expected reward from correct and incorrect responses on each trial. We found that unequal reward caused similar shifts in target selection (reward bias) between the two experiments. Moreover, observed reward biases were independent of the individual’s sensitivity to sensory signals. These findings suggest that the observed reward effects were determined heuristically via modulation of decision-making processes instead of sensory processing and thus, are more compatible with response bias rather than perceptual bias. To further explain our findings and uncover plausible neural mechanisms, we simulated our experiments with a cortical network model and tested alternative mechanisms for how reward could exert its influence. We found that our observations are more compatible with reward-dependent input to the output layer of the decision circuit. Together, our results suggest that during a temporal judgment task, the influence of reward information on perceptual choice is more compatible with changing later stages of decision making rather than early sensory processing.

## Introduction

Imagine deciding whether your iPhone or your friend’s Google Pixel takes sharper photos. To make this decision impartially, you look at some photos on both phones but end up favoring your own phone. Your decision could be entirely based on perceived quality of photos on the two phones but also could be influenced by fun memories evoked by photos on your phone. Likewise, any perceptual decision making could depend not only on sensory evidence favoring alternative outcomes but also on experienced or expected reward (Sugrue, Corrado, & Newsome, 2005). Understanding how additional information, such as reward, is incorporated into perceptual choice can provide valuable insights into the neural mechanisms underlying of both decision-making and reward processes (Christopoulos, Bonaiuto, & Andersen, 2015; Christopoulos & Schrater, 2015; Farashahi, Ting, Kao, Wu, & Soltani, 2018; Gao, Tortell, & McClelland, 2011; Rorie, Gao, McClelland, & Newsome, 2010; Stanford, Shankar, Massoglia, Costello, & Salinas, 2010; Sugrue et al., 2005).

Recently, there have been a number of studies that investigated the mechanisms by which reward information influences perceptual decision making (Cicmil, Cumming, Parker, & Krug, 2015; Diederich, 2008; Diederich & Busemeyer, 2006; Farashahi, Ting, et al., 2018; Gao et al., 2011; Liston & Stone, 2008; Rajsic, Perera, & Pratt, 2017; Tosoni, Committeri, Calluso, & Galati, 2017; Voss, Rothermund, & Brandtstädter, 2008). Some of these studies suggest that reward information mainly affects perceptual choice by altering the starting or end point of decisionmaking processes (Diederich, 2008; Diederich & Busemeyer, 2006; Feng, Holmes, Rorie, & Newsome, 2009; Gao et al., 2011; Mulder, Wagenmakers, Ratcliff, Boekel, & Forstmann, 2012; Rorie et al., 2010; Summerfield & Koechlin, 2010). For example, Diederich and colleagues propose that value-based perceptual decision making follows a two-stage process in which the payoff of the alternative choices are evaluated first before sensory information is processed independently of the payoff information (Diederich, 2008; Diederich & Busemeyer, 2006). Others, however, argue that reward directly influences the processing of sensory information and perception (Cicmil et al., 2015; Liston & Stone, 2008; Pleger, Blankenburg, Ruff, Driver, & Dolan, 2008; Voss et al., 2008).

These competing hypotheses —influence of reward on sensory processing or influence on later stages of decision making —predict that unequal expected reward should result in perceptual bias or response bias, respectively (Liston & Stone, 2008). That is, by changing sensory processing, reward could alter perception (perceptual bias) similarly to the effect of selective attention on contrast judgment (Carrasco & Barbot, 2019; Carrasco, Ling, & Read, 2004). Going back to our phone analogy, reward could make the more rewarding photos to appear sharper. On the other hand, modulation of later decision processes could bias response toward the more rewarding option (response bias) without any changes in perception. In our analogy, this corresponds to favoring our phone without perceiving any difference in image quality. Nonetheless, evidence supporting either hypothesis mostly has been based on fitting performance and reaction time data using different models (e.g., drift diffusion model) and thus, is model dependent. Moreover, it is unclear whether positive and negative expected reward outcomes influence perceptual decision making similarly or differently.

To address these questions and distinguish between the two alternative hypotheses, we used two sets of experiments to directly measure the influence of unequal expected reward on perception and on choice during a temporal judgment task. More specifically, in Experiments 1 and 2, subjects made saccades to report what they perceived to be the first or second of the two identical targets that appeared on the computer screen with varying onset asynchrony. We manipulated the amount of reward points to be gained (gains) or lost (losses) upon correct and incorrect response, respectively, on each trial. Different values of expected gains and losses associated with the left and right choices were presented on the two sides of the fixation cross to create three reward conditions: neutral, gain, and loss. This design allowed us to estimate the shift in target selection and sensitivity of choice in response to different target onset asynchronies (TOAs) and how this shift and sensitivity were affected by reward manipulation.

In both experiments, reward information was not predictive of the correct or incorrect response on a given trial and thus, could be entirely ignored (i.e., subjects only needed to attend to the timing of the two targets in order to perform the task), causing no bias in choice or perception. However, reward information could bias the subject to select the target on the more rewarding side when detecting the first or second target was difficult (response bias). Because of its salience, reward information could also shift attention to the more rewarding side and facilitate target detection on that side (perceptual bias). By comparing reward-induced shifts in target selection during Experiments 1 and 2, we determined whether reward caused perceptual bias and/or response bias. We expected that a perceptual bias would result in opposite shifts in target selection, whereas a response bias would cause similar shifts in the two experiments (see Results for more details). Finally, we used a biophysically-plausible cortical network model to replicate the experimental data in order to identify possible neural mechanisms underlying the influence of reward on perceptual decision making.

## Materials and Methods

### Ethics statement

A total of 29 (15 female) subjects were recruited from the Dartmouth College student population (ages 18–22 years) to participate in our experiments. Of the 29 subjects, 21 subjects performed in both Experiments 1 and 2, each of which consisted of 4 sessions. The remaining 8 subjects only performed in either Experiment 1 or 2. All participants gave informed consent to participate according to a protocol approved by the Dartmouth College Institutional Review Board. All subjects signed a written consent form before participating in the experiments.

### Experimental design

This study used a within-subject design in which human subjects performed two variations of a temporal judgment task known as the paired-target task under three different reward conditions (neutral, gain, and loss; see *Reward Conditions*). The subjects were required to detect the order of two visually-identical targets (gabor patches) that appeared on the computer screen at varying time intervals with respect to each other (Schiller & Chou, 1998). The subject’s task was to saccade to the first target in Experiment 1 (after both targets were presented) whereas in Experiment 2, the subject’s task was to saccade to the second target (**Fig. 1**). Subjects were instructed to saccade to the correct target (first target in Experiment 1 and second target in Experiment 2) without being concerned about the speed (i.e., reward did not depend on response time). Following each response, subjects received reward feedback – a green or red circle with the amount of reward points earned or lost, respectively (**Fig. 1**).

**Figure 1.**
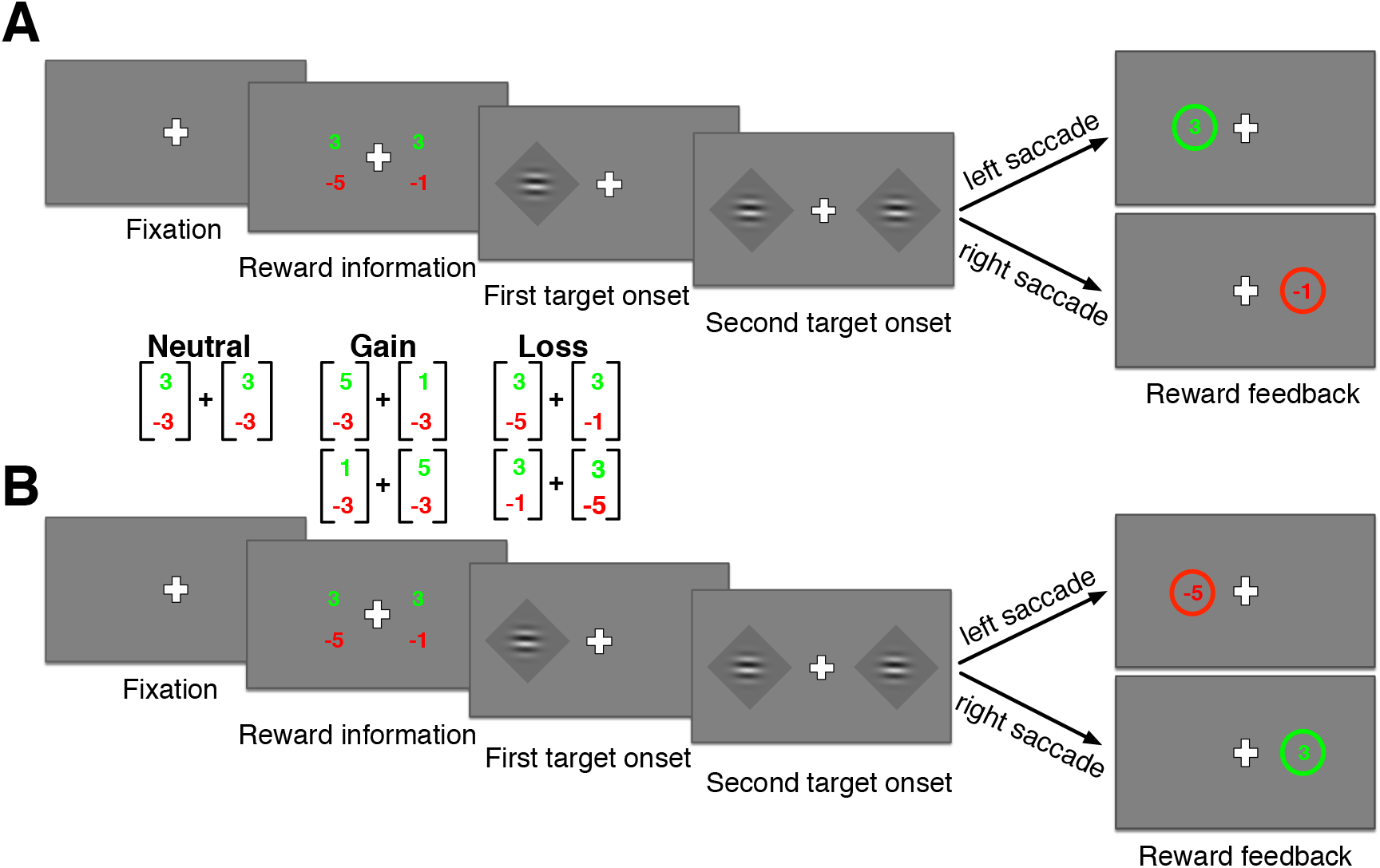
Schematic of the experimental paradigm. (**A**) Timeline of a trial during Experiment 1. Each trial began with a fixation cross followed by the presentation of reward points associated with a correct/incorrect response for each target on both sides of the fixation cross (in green for gains and red for losses). The amount of expected reward was manipulated in three experimental conditions (neutral, gain, and loss) as indicated in the inset. Following the presentation of the reward information, two identical targets (gabor patches) appeared on the screen asynchronously. The subjects’ task was to report the first target that appeared on the screen by making a saccade to the target. Reward feedback was then given by a green circle for correct response or red circle for incorrect response around the selected choice with reward points gained or lost, respectively, in the center of the circle. (**B**) Timeline of a trial during Experiment 2. Stimulus and reward information were presented similar to Experiment 1 except the correct response required making a saccade to the second target that appeared on the screen.

Subjects total reward points were exchanged for a monetary reward at the end of each experiment based on their performance. They were compensated with a combination of money and “t-points,” extra-credit points for classes within the department of Psychological and Brain Sciences at Dartmouth College. More specifically, in addition to the base rate of $10/hour or 1 t-point/hour, subjects were compensated up to an additional $10/hour depending on the total reward points they collected in each experiment. The order of two experiments was randomized across subjects to avoid possible confounds.

At the beginning of each trial, subjects were required to fixate on a white cross at the center of the screen for at least 500 ms (see **Fig. 1** for the trial sequence). Fixation was considered broken when the subject’s eye position deviated 112.5 pixels (~2° visual angle) from the fixation cross before the fixation period ended. If the subject broke fixation, a new trial would begin after a 1000 ms pause. After successful fixation, the amounts of reward points to be earned/lost upon a correct/incorrect response to each target were signaled on the side of the fixation cross corresponding to that target. The amounts of reward points expected from correct and incorrect responses, which we refer to as gains and losses, on both sides were presented close to the fixation cross and large enough to be read without breaking fixation (~2 ° visual angle from the fixation cross). To allow more sensitivity to reward information, the amounts of gains and losses were presented in green and red, respectively. After the offset of reward information at ~1000 ms, there was a variable interval between 500 to 1000 ms (uniform distribution) before the first target appeared on the screen. The second target then appeared after an interval selected from the following values: 0, 16.7, 33.3, 50, and 66.7 ms. We refer to these values as the target onset asynchrony (TOA). Targets were presented at equal distances from the fixation cross (~ 7° visual angle).

All stimuli were presented on an FSI AM250 monitor, which has a refresh rate of 60 Hz and resolution of 1920 × 1080 pixels. Subjects were seated 60 cm from the computer screen. Eye movements were recorded using a video-based eye-tracking system (Eyelink 1000, SR Research Ltd, Mississauga, Canada). To minimize head movements, subjects were seated with their chin on a chin rest. The experiments were programmed using PsychToolbox in MATLAB (Brainard, 1997; Kleiner et al., 2007; Pelli, 1997).

### Reward conditions

Both Experiments 1 and 2 consisted of four sessions, each of which corresponded with one of the three reward conditions: neutral, gain, and loss. In the neutral condition, which always preceded either the gain or loss condition, correct and incorrect responses to either target resulted in gaining or losing 3 points, respectively (indicated by the [3,-3] on the two sides of the fixation point; **Fig. 1**). In the gain condition, reward points for saccade to one target were [5,-3], whereas the other target had reward points of [1,-3]. In the loss condition, the targets were associated with reward points of [3,-1] and [3,-5]. These values were selected based on our pilot study to ensure similar differences in gain and loss values, assuming an average loss-aversion factor of 2. During the gain and loss conditions, the side with the target with higher expected value was randomly assigned on each trial. The order of gain and loss conditions was randomized across subjects. Each reward condition (gain, loss, or neutral) consists of 180 trials and was performed in a single session of the experiment without pause (lasting about 15-20 minutes). Therefore, each session corresponded to one of the three reward conditions.

### Fitting choice behavior and estimated parameters

We fit choice data from each session (reward condition) of the experiment separately to estimate different aspects of target selection. The psychometric function in Experiment 1 was defined as the probability of choosing the left target as a function of the TOA favoring the left target, TOA = *t*_right_ - *t*_left_, where *t*_left_ and *t*_right_ are the onset time of the left and right targets, respectively. The psychometric function in Experiment 2 was defined as the probability of choosing the left target as a function of the TOA favoring the right target (by flipping the sign for the TOA on each trial) in order to measure target selection as a function of the signal relevant for the task. We then used the standard maximum likelihood estimation (by minimizing the negative log likelihood) to fit the psychometric function in each session using the following equation:

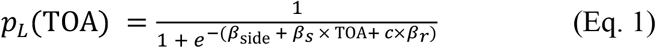

where *β_side_* indicates the subject’s preference for saccade to target on the left side of the fixation cross regardless of what was presented, *β_s_* represents sensitivity to sensory information (i.e., how selection changes as a function of the TOA), *β_r_* denotes reward bias (i.e., preference for saccade to the target with the larger expected reward). Finally, *c* is a dummy variable that indicates the more rewarding side on each trial (i.e., it is 1 if the left side is more rewarding, 0 if both sides are equally rewarding, and −1 if the right side is more rewarding). Critically, these parameters capture difference aspects of target selection and how it is influenced by reward information.

We also computed error in parameter estimation with two different methods. Firstly, we used the Hessian matrix of the log-likelihood function to find the 95% confidence interval of the estimated parameters. Secondly, we randomly sampled 95% of the data 50 times to fit the ensuing psychometric function and then calculated the mean and standard deviation of the estimated parameters across all samples.

### Data exclusion

We excluded 20 sessions from the total 200 sessions of the two experiments completed by the 29 subjects. Exclusion was performed on a session-by-session basis using three exclusion criteria. First, we excluded sessions in which sensitivity to the TOA was negative, indicating that the subject did not perform the task properly by ignoring the main task variable (TOA) on most trials. Second, we excluded sessions in which the overall task performance did not exceed chance (50%) plus two times s.e.m., reflecting an unusually poor performance. Using these two criteria, we removed 19 sessions from eight different subjects (seven of these subjects had 1 or 2 excluded sessions except one subject whose all 8 sessions were excluded). Finally, we discarded sessions in which either of the fitting parameters (i.e., *β_s_* (sensitivity), *β_side_*, or *β_r_*) deviated by more than three times the standard deviation from the corresponding parameter’s mean across all sessions. The third criteria led us to remove 1 more session. Results reported here are based on the remaining 180 sessions (valid sessions).

### Statistical and data analysis

We used the standard maximum likelihood estimation to find the parameters of the psychometric function in each session. Then, we used two-sided signed rank test to compare the estimated parameters with the null hypothesis (0 corresponding to no effect). To compare the reward biases of the loss and gain conditions, we used two-sided Wilcoxon ranksum test. We used both Pearson and Spearman correlation to examine the correlation between estimated parameters.

To determine the model that best explains the variances in the saccadic reaction time, we used a stepwise general regression model (GLM). We included the following regressors in the stepwise GLM: unequal reward condition indicating unequal (gain and loss) or equal (neutral) expected reward outcomes, the TOA, response accuracy (correct vs. incorrect response), and a dummy variable indicating whether the chosen target had the higher or lower expected reward (chosen-target relative value). The last regressor only applied to the gain and loss conditions. A stepwise GLM procedure examines all combinations of regressors and their interactions to determine terms whose inclusion results in a significant increase of the adjusted-*R^2^*. We used custom codes and the statistical package in MATLAB (MathWorks Inc., Natick, MA) to perform all simulations and statistical analyses.

### Optimality analysis

The optimal reward bias is defined as the amount of shift in target selection that maximizes the total reward earned in a given session. To determine the optimal reward bias, we first calculated the expected amount of reward earned assuming a given level of sensitivity to sensory information and loss aversion. The subject’s sensitivity can predict the overall number of correct choices and thus, the number of reward points they can earn. Loss aversion causes individuals to respond more strongly to reward points lost than gained (note that reward points were assigned to the two sides on a trial-by-trial basis). Considering these factors, the expected payoff associated with saccades to the left (*L*) and right (*R*) can be calculated using the following equations:

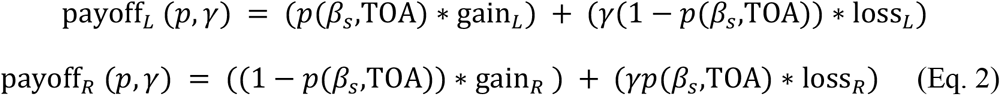

where *γ* represents the loss-aversion factor and *p*(*β_s_,TOA*) is the probability of the correct response for a given TOA:

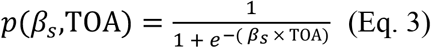

where *β_s_* represents sensitivity to sensory information. The total expected payoff for a given value of shift in target selection is equal to:

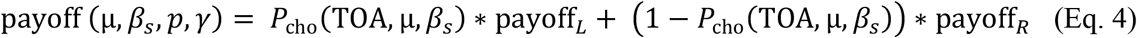

where *μ* is the amount of shift in target selection, and *P*_cho_(TOA, μ, *β_s_*) represents the probability of choice for a given TOA, *μ,* and *β_s_*, which is computed as follows:

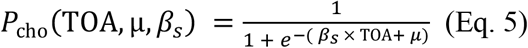

As a result, the total expected amount of reward earned for all values of *p* for given values of *μ* and *β_s_* is equal to

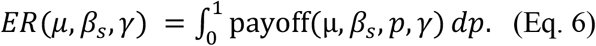

where the integral is computed by binning and summating over all values of *p.* Finally, the optimal reward bias can be computed by finding a value of shift in target selection that maximizes the value of *ER* for specific values of sensitivity and loss-aversion factor (using Eq. 6). We used predicted values of optimal reward bias to examine reward bias exhibited by individual subjects.

### Computational model

The basic model consisted of two cortical columns with two pools of excitatory neurons and one inhibitory pool of neurons in the superficial layer and two pools of excitatory neurons in the deep layer (Soltani, Noudoost, & Moore, 2013) (**Supplementary Fig. 1**). All neural pools in all the layers received a background input mimicking input from adjacent cortical neurons with different types of selectivity. The excitatory pools in the superficial layer also received visual input related to the presentation of targets on the screen. Specifically, the visual input to the two pools were similar except that they had different onset timing according to the TOA on each trial. Moreover, the two pools of excitatory neurons in the superficial layer were mutually inhibited using a shared pool of inhibitory interneurons. This mutual inhibition created a winner-take-all competition and caused activity in excitatory pools to diverge in both the superficial and deep layers. We used a mean-field approximation of a spiking network model of decision making to simulate the superficial layer (Wong & Wang, 2006). Each excitatory pool of neurons in the deep layer had weak self-excitatory recurrent connections. The deep layer then projected its output to the brain stem or superior colliculus to direct a saccadic eye movement. We determined the choice of the network on each trial by identifying the first deep-layer excitatory pool whose activity passed 15 Hz (considered the winner pool). The full details of the basic model are described elsewhere (Soltani et al., 2013).

To simulate the observed effects of unequal reward information, we considered three alternative mechanisms. These mechanisms affected different parts of the model, mimicking either modulations of sensory processing or later stages of decision-making processes. First, to simulate the effect of reward information on later stages of decision-making processes, we included a reward-based input to the excitatory pools in the deep layer (**Fig. 9A**; Mechanism 1). This input was independent of the amount and timing of the visual input (TOA) on a given trial. Second, to simulate the effect of reward information on sensory processing, we assumed two alternative mechanisms in which reward information could modulate the visual input to the decision circuit. In the first mechanism, the input evoked by targets with a larger and smaller expected reward was multiplied by (1 + λ) and (1 – λ), respectively, where *λ* is a constant that measures the modulation of visual input by reward information (**Fig. 9B**; Mechanism 2). This results in stronger input for the target on the more rewarding side. In the second mechanism for modulating sensory processing, the input for the target with larger expected reward was modulated through a shift in TOA in favor of the target with larger expected reward (**Fig. 9B**; Mechanism 3). This shift mimics faster processing of input related to the target on the more rewarding side in higher visual areas. In both Mechanisms 2 and 3, the reward cue presented to subjects modulates the processing of sensory information and thus, could result in perceptual bias. Finally, in order to simulate Experiment 2 (selecting the second target that appears on the screen), we assumed that the projections of the output in excitatory pools of the superficial layer is switched via a gating mechanism (green dashed lines in **Fig. 9A, B**) to allow the selection of the non-winner pool as the response.

Due to the nonlinear dynamics of the proposed network models, we had no *a priori* predictions about which alternative mechanism would be more compatible with the observed lack of correlation between shifts in target selection and the sensitivity to the TOA, and whether a single mechanism was sufficient to capture our main experimental findings. Also, the two proposed mechanisms of reward influence on sensory processing – Mechanism 2 corresponding to enhancement and Mechanism 3 to facilitation of sensory signals due to unequal reward – were different enough to warrant the simulation and examination of both mechanisms.

## Results

### Alternative mechanisms for the influence of reward on target selection and their predictions

To study the influence of reward information on perceptual decision making in general and in temporal judgment in particular, we used modified versions of the paired-target task in which the subjects reported what they perceived to be the first (Experiment 1) and second (Experiment 2) of the two targets to appear on the screen by making a saccade to the target. At the beginning of each trial, the amounts of reward points expected to be gained and lost upon correct and incorrect responses, respectively, were presented on the two sides of the fixation cross and were manipulated across experimental sessions (**Fig. 1**; see Materials and Methods).

From the point of view of an ideal observer, reward information could be used to enhance the overall sensitivity to sensory information without inducing any bias (perceptual or response). This could happen because in our experiments, the expected reward points associated with selection of the targets on the two sides does not predict the correct response on a given trial. Alternatively, the ideal observer could use reward information to bias its response. For a non-ideal observer, however, expected reward could have a profound effect on both perception and choice. On the one hand, attention could be automatically employed on the more rewarding side, resulting in an enhanced processing of the target on that side (perceptual bias) even though this enhancement does not improve correct detection. On the other hand, because of task difficulty (especially on trials with no or short TOA), subjects could use reward information to increase selection of the target on the more rewarding side (response bias) to improve the overall reward. As explained below, these alternative effects of expected reward could be dissociated based on the results in Experiments 1 and 2.

If reward information results in an enhanced processing of the more rewarding side (i.e., perceptual bias) for a non-ideal observer, the target on the more rewarding side would be perceived to appear earlier when both targets appeared on the screen simultaneously. This would increase the probability of choosing the target on the more rewarding side in Experiment 1 (**Fig. 2A**). In contrast, earlier perception of the target on the more rewarding side would increase the probability of choosing the target on the less rewarding side in Experiment 2 in which the subjects have to saccade to the target that appeared second (**Fig. 2B**). If instead, reward information results in more frequent selection of the target on the more rewarding side without any changes in sensory processing (i.e., response bias), the target on the more rewarding side would be selected with a higher probability in both Experiments 1 and 2 (**Fig. 2C-D**). Therefore, perceptual bias would result in opposite shifts in target selection in Experiments 1 and 2, but response bias would shift target selection (toward the better side) similarly in both experiments.

**Figure 2.**
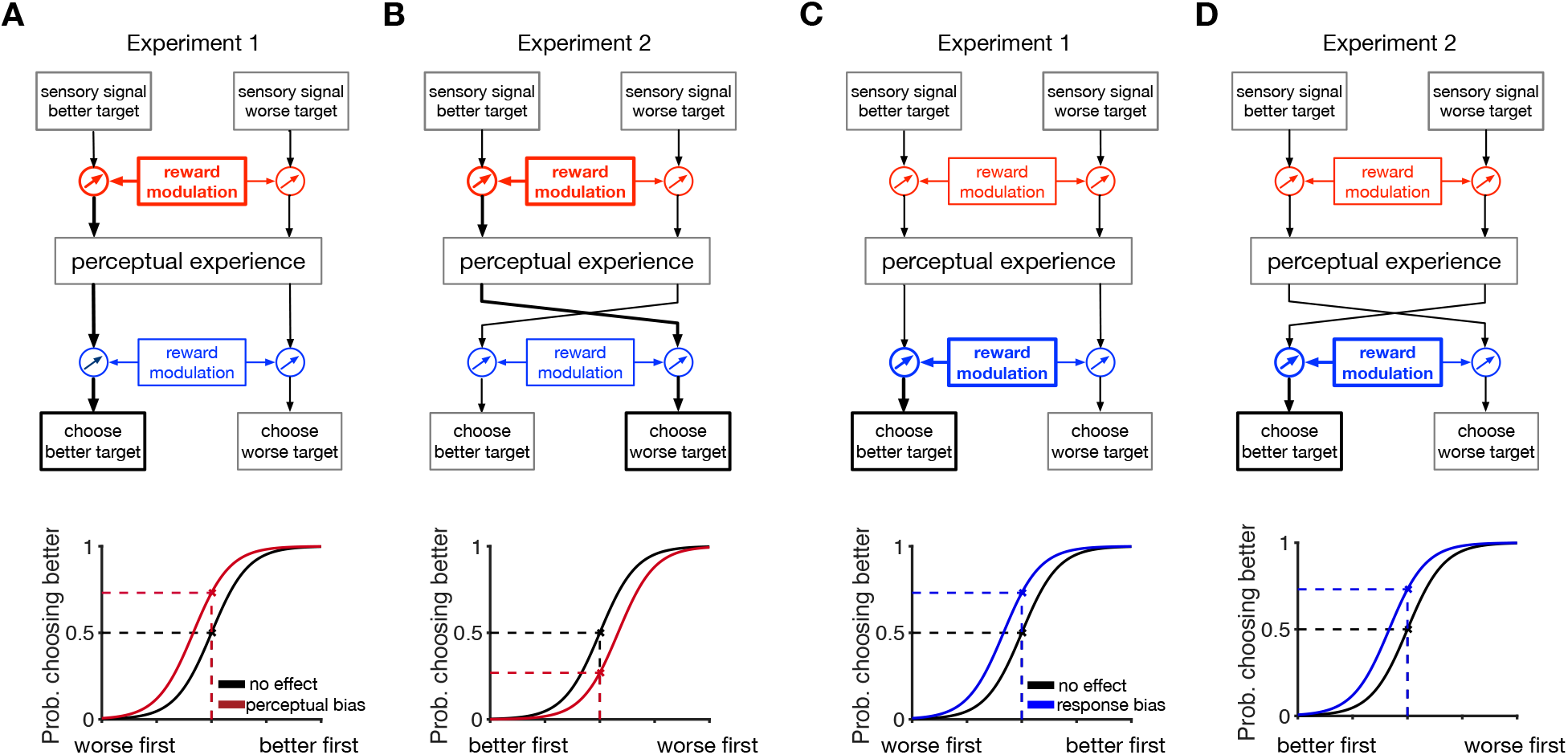
The effects of reward-induced perceptual bias and response bias on target selection in Experiments 1 and 2. **(A-B)** The effect of perceptual bias on target selection in Experiments 1 (A) and 2 (B). Perceptual bias can enhance or facilitate processing of the target that appear on the more rewarding side (depicted with thicker lines) causing this target to be perceived earlier. This would increase the probability of choosing the target on the more rewarding side in Experiment 1 (A) but increase the probability of choosing the target on the less rewarding side in Experiment 2, in which the subjects have to saccade to the target that appeared second (B). The insets at the bottom depict changes in the probability of choosing the better target due to perceptual bias as the function of the TOA favoring the better (worse) target in Experiment 1 (respectively, Experiment 2). The black curve shows the probability of choosing the better target in the absence of any reward modulation. The highlighted arrow indicates the locus of reward modulation. **(C-D)** The effect of response bias on target selection in Experiments 1 (C) and 2 (D). Response bias increases the probability of choosing the better target in both experiments. Conventions are the same as in panels A-B.

In addition to the pattern of a shift in target selection due to unequal expected reward, the relationship between this shift and sensitivity to sensory information can be used to distinguish between alternative scenarios. To establish this relationship, we computed the optimal shift based on given values of sensitivity to sensory evidence (*β_s_*) and loss-aversion factor (*γ*) separately in the gain and loss conditions (see *optimality analysis* in **Materials and Methods**). We found that in the gain condition, the optimal reward bias should decrease as loss aversion increases for a given level of sensitivity (**Fig. 3B**). Moreover, the optimal bias should decrease with larger sensitivity (**Fig. 3C**). In contrast, in the loss condition, the optimal reward bias should increase with more loss aversion (**Fig. 3E**) but decrease with larger sensitivity. These results illustrate that optimal shift in target selection requires that reward bias to be inversely correlated with the individual’s level of sensitivity to sensory evidence. Moreover, loss aversion should have opposite effects on reward bias in the gain and loss conditions.

**Figure 3.**
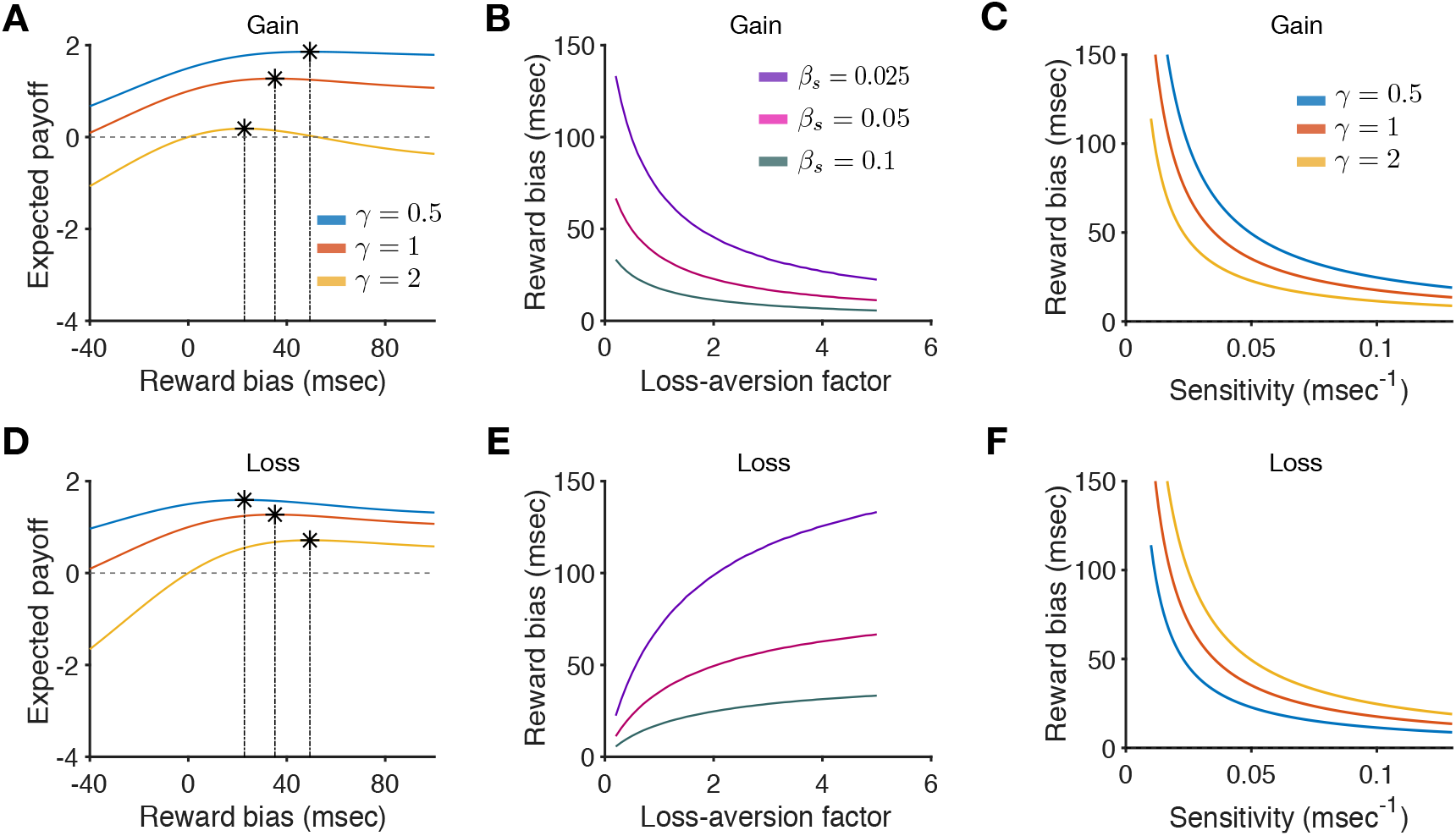
Predicted optimal shift in target selection due to unequal expected reward. (**A**) The expected payoff for a given level of sensitivity (*β_s_* = 0.05 ms^−1^) and reward bias during the gain condition. Each curve represents the expected payoff for a different level of loss aversion (loss-aversion factor *γ*). The peak in each curve indicates the optimal reward bias. (**B**) The optimal reward bias as a function of the loss-aversion factor in the gain condition separately for three different values of the sensitivity. The optimal reward bias decreases as loss aversion increases in the gain condition. (**C**) The optimal reward bias as a function of the sensitivity to sensory evidence. The optimal reward bias diminishes as the sensitivity increases. (**D-F**) The same as in panels A-C but for the loss condition. The optimal reward bias increases as loss aversion increases in the loss condition. Nevertheless, the optimal reward bias decreases with larger values of the sensitivity similar to the gain condition.

Together, these results predict that perceptual bias would result in opposite shifts in target selection in Experiments 1 and 2 (**Fig. 3A**), but response bias would shift target selection (toward the better side) in the two experiments similarly (**Fig. 3B**). In contrast, an ideal observer may exhibit no bias or equal bias in the two experiments by changing the decision criteria based on reward information (Gold & Shadlen, 2001) (**Fig. 3C**). Additionally, in case of perceptual bias due to asymmetric changes in sensory processing or response bias adopted by an ideal observer, an optimal shift in target selection depends on sensitivity to sensory information; the shift should be small when sensitivity is high (corresponding to good temporal judgment) and large if sensitivity is low, corresponding to poor temporal judgment (**Fig. 3D, F**). On the other hand, shift in target selection could be independent of sensitivity to sensory information if unequal expected reward causes response bias (**Fig. 3E**).

### Influence of reward information on target selection

We then used our experimental data to test alternative predictions about the effects of reward on target selection depicted in **Fig. 4**. To that end, we fit each subject’s psychometric function (the probability of choosing the left target as a function of the TOA) using a sigmoid function in order to estimate three parameters for each subject: sensitivity to sensory information (*β_s_*), side bias (*β*_side_), and reward bias (*β_r_*) (see Materials and Methods for more details). These parameters capture three features of target selection in our experiment: sensitivity measures the fidelity of target selection to sensory evidence (i.e., TOA), side bias measures an overall bias in choosing the left or right target independently of sensory information, and reward bias measures the shift in target selection toward the more rewarding side (**Fig. 5A-C**). These parameters were estimated separately for each of the three reward conditions. To minimize possible side bias, the side associated with the target with better reward outcomes had been randomly assigned for each trial.

**Figure 4.**
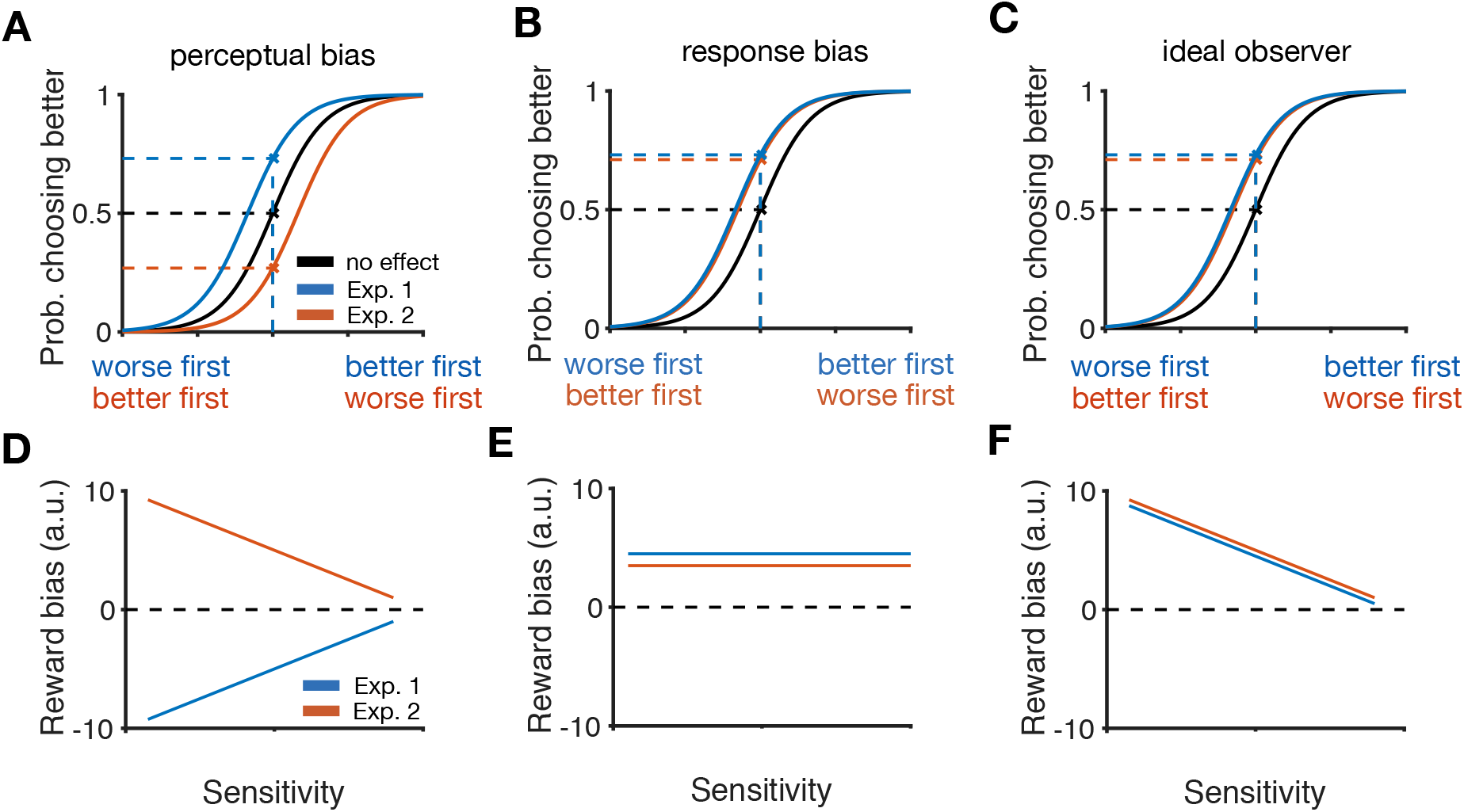
Reward bias and its relationship to sensitivity to sensory information as predicted by different mechanisms for the influence of reward. **(A)** Perceptual bias caused by differential processing of the two targets results in opposite shifts in target selection in the two experiment. Plot depicts a hypothetical psychometric function for a non-ideal observer. The blue and orange labels for the x-axis are corresponding to the Experiments 1 and 2, respectively. Conventions are the same as in Figure 2. **(B)** A non-ideal observer with response bias exhibit similar shifts in target selection in the two experiments. **(C)** The effect of reward on target selection for an ideal observer. **(D-E)** Prediction of correlation between reward bias (shifts in target selection due to reward) and sensitivity to sensory information for a non-ideal observer exhibiting perceptual (D) or response bias (E). **(F)** Prediction of correlation between reward bias and sensitivity for an ideal observer, separately for each experiment. Using the same mechanism for shift in target selection results in similar reward bias in the two experiments.

**Figure 5.**
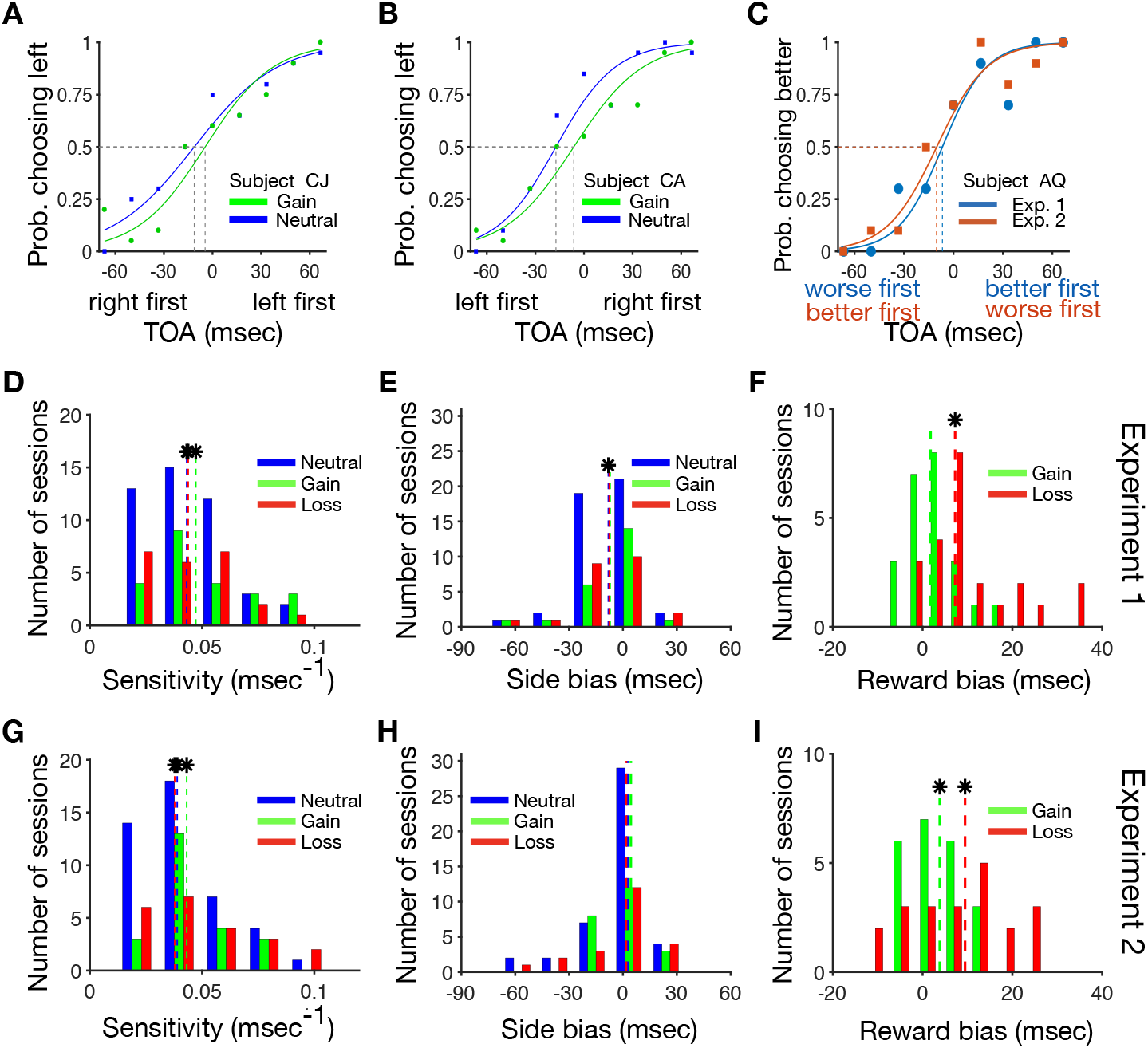
Influences of unequal expected reward on target selection. (**A-C**) Example psychometric functions in different conditions of Experiments 1 and 2. (**A**) Plotted is the probability of choosing the left target as a function of the TOA for an example subject in the gain and neutral conditions of Experiment 1. This subject exhibited a slightly larger level of sensitivity in the gain compared to the neutral condition. Positive (negative) values of TOA correspond to the left (right) target to appear first. (**B**) The same as in panel A but for another subject who exhibited a larger side bias in the neutral than in the gain condition of Experiment 2. In these plots, side bias is equal to the TOA at which the probability of choosing the two targets are equal. In Experiment 2, negative (positive) values of TOA correspond to the left (right) target appearing first. (**C**) Psychometric functions of an example subject in the gain condition of Experiments 1 and 2. Plotted is the probability of choosing the better target as a function of the TOA for that target. For Experiment 1, negative (positive) values of TOA means the target with smaller (larger) expected reward has appeared first. In Experiment 2, negative (positive) values of TOA means the target with smaller (larger) expected reward appeared second. In these plots, reward bias can be interpreted as the TOA at which the subject chooses the two targets equally. This subject exhibited similar reward bias in the two experiments. (**D-F**) Histograms plot the number of subjects with given values of sensitivity (**D**), side bias (**E**), and reward bias (**F**) under different reward conditions during Experiment 1. The dashed lines show the medians, and each asterisk indicates a significant difference from 0 (two-sided signed-rank test, *p* < 0.05). (**G-I**) The same as in panels D-F but for Experiment 2.

We found that subjects were sensitive to the TOA in both experiments. The average values (±std) of sensitivity in Experiment 1 were equal to: 0.0432±0.0191, 0.0472±0.0222, and 0.0438±0.0179 (ms^-1^) for the neutral, gain, and loss conditions, respectively. Notably, these average values were larger than zero (two-sided signed-rank test; neutral: *p* = 5.18 × 10^−9^, *d* = 2.25; gain: *p* = 27 × 10^−5^, *d* = 2.29; loss: *p* = 2.7 × 10^−5^, *d* = 2.47; **Fig. 5D**). The average values of sensitivity in Experiment 2 were equal to: 0.0389±0.0195, 0.0432±0.0187, and 0.0378±0.0259 (ms^−1^) for the neutral, gain, and loss conditions, respectively. As in Experiment 1, these average values were larger than zero (two-sided signed-rank test; neutral: *p* = 7.6 × 10^−9^, *d* = 2.08; gain: *p* = 2.7 × 10^−5^, *d* = 2.34; loss: *p* = 4 × 10^−5^, *d* = 1.76; **Fig. 5G**).

We also examined estimated side bias in different reward conditions and experiments. In Experiment 1, we did not observe any evidence for side bias except in the neutral condition (twosided signed-rank test; neutral: −8.11±17.21 ms, *p* = 0.002, *d* = 0.47; gain: −7.14 ±16.60 ms, *p* = 0.083, *d* = 0.40; loss: −7.60±22.72 ms, *p* = 0.083, *d* = 0.36; **Fig. 5E**). Furthermore, we did not find any evidence for side bias in any conditions of Experiment 2 (two-sided signed-rank test; neutral: 2.54±19.77 ms, *p* = 0.55, *d* = 0.12; gain: 4.41±14.63 ms, *p* = 0.72, *d* = 0.06; loss: 1.69±22.20 ms, *p* = 0.88, *d* = 0.13; **Fig. 5H**). Together, these results illustrate that subjects exhibited very small side bias in both experiments.

We then examined reward bias or the shift in target selection due to unequal expected reward. In Experiment 1, 87% and 96% of subjects exhibited a significant reward bias in the gain and loss conditions, respectively. However, across all subjects, reward bias toward the more rewarding side was significant only in the loss condition (two-sided signed-rank test; gain: 1.79±5.92 ms, *p* = 0.128, *d* = 0.36; loss: 7.25±10.56 ms, *p* = 0.0001, *d* = 1.0; **Fig. 5F**). Furthermore, this shift in target selection was stronger during the loss than gain condition (two-sided Wilcoxon ranksum test; Δ = 5.45 ms, *p* = 0.002, *d* = 0.7). In Experiment 2, 83% and 96% of subjects exhibited a significant reward bias in the gain and loss conditions, respectively, and across all subjects, reward bias were significantly larger than zero in both gain and loss conditions (twosided signed-rank test; gain: 3.82±10.40 ms, *p* = 0.041, *d* = 0.43; loss: 9.45±14.00, ms, *p* = 0.006, *d* = 0.67; **Fig. 5I**).

Finally, to ascertain that noise in the estimation of the parameters did not influence our results, we calculated the error of estimation based on two different methods (see Materials and Methods for more details). Using the first method, the Hessian matrix of the log-likelihood function, we found small errors (mean ~8%) in the estimation of reward biases (**Supplementary Fig. 2**). The second method that was based on resampling produced a slightly larger error estimation (mean ~24%; **Supplementary Fig. 3**). Therefore, the errors calculated by either method do not invalidate our main conclusions and demonstrate the robustness of our fitting procedure.

We also assessed the relationship between reward bias in the gain and loss conditions within each individual subject. However, we did not find any evidence for correlation between reward bias in the gain and loss condition for subjects who successfully performed (valid sessions) in both conditions (Pearson correlation; Experiment 1: *r* = –0.24, *p* = 0.32, *N* = 20; Experiment 2: *r* = 0.09, *p* = 0.72, *N* = 17; Spearman correlation; Experiment 1: *r* = –0.17, *p* = 0.46, *N* = 20; Experiment 2: *r* = –0.11, *p* = 0.67, *N* = 17). Combining data from both experiments, we observed an overall larger bias in the loss than in the gain condition (two-sided signed-rank test; Δ = 6.00±12.73, *p* = 0.0085, *d* = 0.48, *N* = 37; **Fig. 6**). This observed bias provides evidence for loss aversion in perceptual decision making with possible gains and losses.

**Figure 6.**
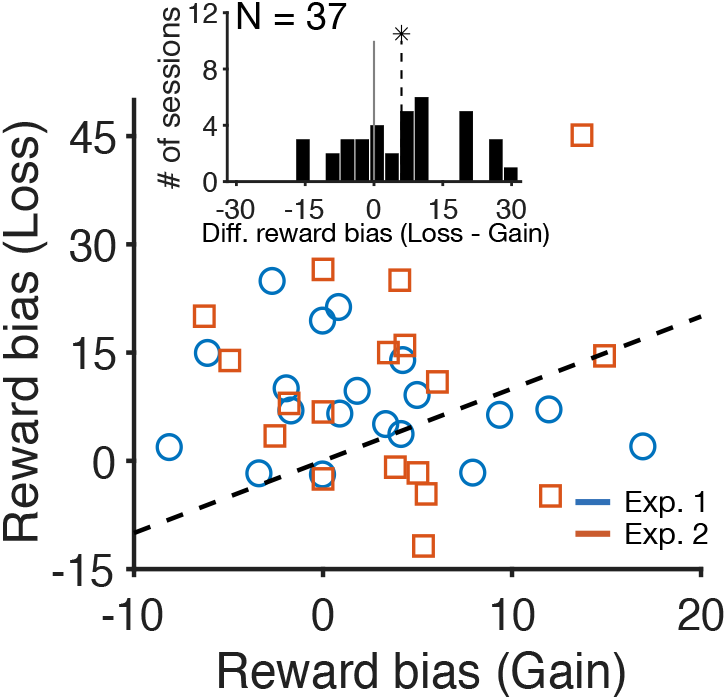
Larger shift in target selection during the loss than in the gain condition. Plotted is individual subjects’ reward bias during the loss condition as a function of reward bias during the gain condition for Experiments 1 (blue) and 2 (red). Here, we only included subjects who performed both experiments successfully (valid sessions). The inset plots the distribution of the difference between reward bias in the loss and gain conditions over both experiments. The dashed line is the median and the asterisk indicates a significant difference from 0 (two-sided signed-rank test, *p* < 0.05).

As discussed earlier, a non-ideal observer could exhibit a shift in target selection due to separate (and not exclusive) mechanisms: differential enhancement or facilitation of sensory processing, which can result in perceptual bias; and/or increase in the tendency to make a saccade to the target on the better side (i.e., response bias). Importantly, these mechanisms predict different patterns of results for Experiments 1 and 2; perceptual bias results in opposite shifts in target selection in the two experiments, whereas a response bias would cause similar shifts in the two experiments (**Fig. 4**). Hence, a difference between shifts in target selection in Experiments 1 and 2 would mostly reflect perceptual bias. Instead, we found no evidence for a difference in reward bias between the two experiments in the gain or in the loss condition (two-sided signed-rank test; gain: Δ = −1.27±13.20 ms, *p* = 0.29, *d* = 0.26; loss: Δ = 2.24±18.79 ms, *p* = 0.74, *d* = 0.02). This result provides evidence more compatible with response bias or an increase in the tendency to make a saccade to the target on the better side.

As mentioned earlier, unequal expected reward could result in response bias for both ideal and non-ideal observers. For a non-ideal observer, response bias is determined heuristically. For an ideal observer, however, response bias could happen due to a shift in decision criteria, making the shift in target selection correlated with sensitivity to sensory information (**Fig. 4**). Therefore, we calculated the correlation between reward biases and sensitivity of the subjects in both the gain and loss conditions of the two experiments. We did not find any evidence for correlation between response bias and an individual’s sensitivity to sensory information (Pearson correlation; gain condition: Experiment 1: *r* = –0.13, *p* = 0.55; Experiment 2: *r* = – 0.34, *p* = 0.11; loss condition: Experiment 1: *r* = –0.17, *p* = 0.44; Experiment 2: *r* = –0.1, *p* = 0.64). This result suggests that the observed shift due to the reward information is more compatible with the response bias in a non-ideal observer.

### Comparison of subjects’ reward bias with optimal values

Our experimental results are more compatible with response bias that was determined heuristically. Nonetheless, we compared the observed and optimal values of shift in target selection due to unequal reward (reward bias) for individual subjects assuming different values of loss aversion (because we did not have access to the degree of loss aversion in individual subjects; **Fig. 3**). We found that in the gain condition of both experiments, subjects exhibited reward biases that were smaller than the predicted optimal biases based on loss neutrality (loss-aversion factor equal to 1) (two-sided signed-rank test; Δ = - 39.58 ± 3.64 ms, *p* = 3.5 × 10^−9^, *d* = 1.76; **Fig. 7A-C**). Therefore, the observed reward biases would be optimal if loss-aversion factor was very large (loss-aversive) because larger loss aversion gives rise to smaller shifts in the gain condition.

**Figure 7.**
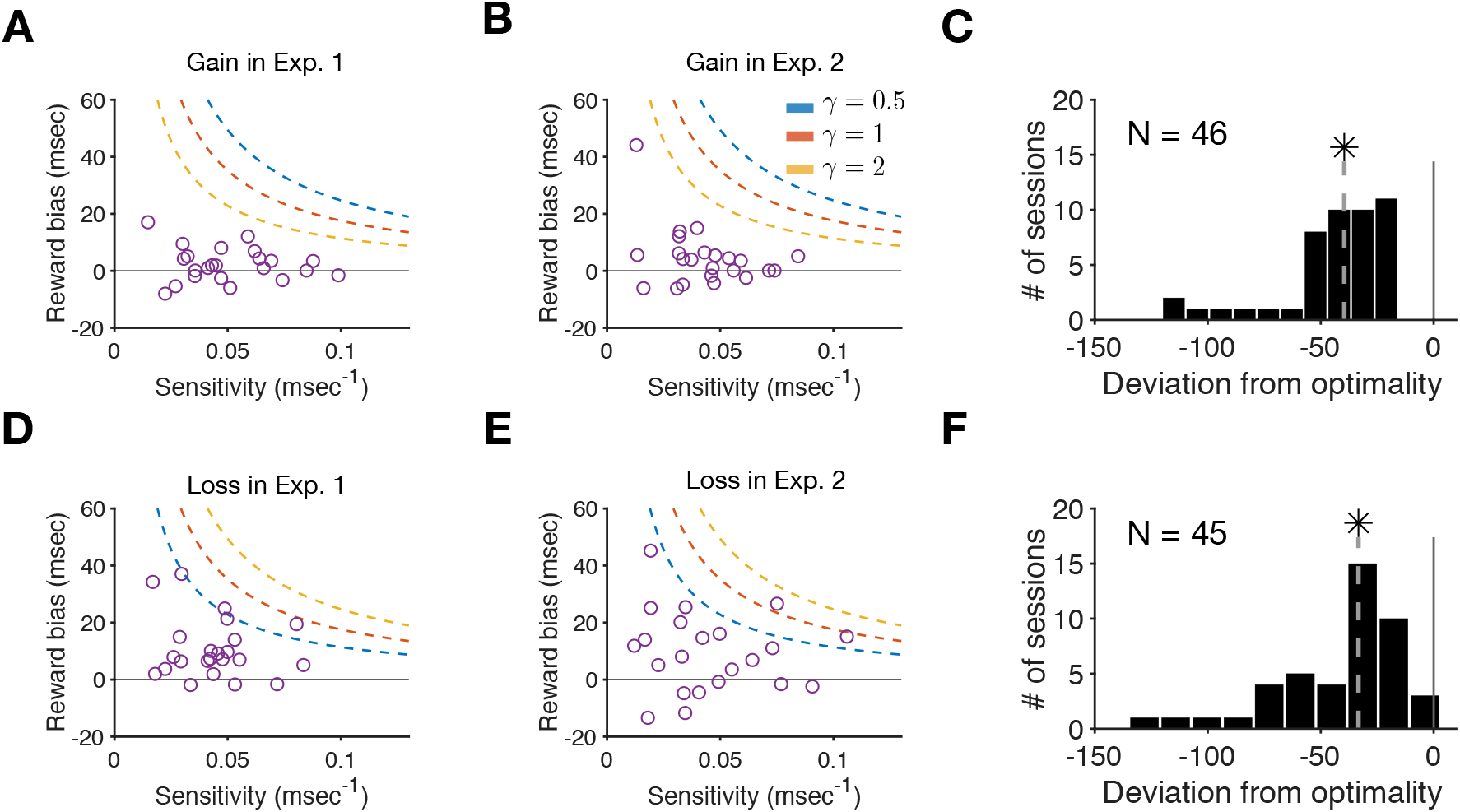
The amount of shift in target selection was suboptimal in the gain and loss conditions of both experiments. (**A**) Plot shows individuals’ reward bias as a function of their sensitivity to the TOA. The blue, red, and yellow lines represent the optimal value of reward bias for different values of loss-aversion factor. (**B**) The same as in panel A but for Experiment 2. (**C**) Plotted is the distribution of the differences between observed and predicted optimal reward biases based on loss neutrality (*γ* = 1) across all subjects in the gain condition of Experiments 1 and 2. (**D-F**) The same as in panels A-C but for the loss condition in the two experiments. Note that a larger loss-aversion factor predicts larger reward bias in the loss condition (the opposite is true for the gain condition).

The observed reward biases in the loss condition were also smaller than the optimal values based on loss neutrality (two-sided signed-rank test; Δ = −33.29 ± 4.29 ms, *p* = 6.3 × 10^−9^, *d* = 1.41; **Fig. 7D-F**). In this case, however, the observed reward biases would be optimal if loss-aversion factor was very small because smaller loss aversion gives rise to smaller shifts in the loss condition. Therefore, the observed smaller-than-optimal shifts in the loss condition point to lossseeking as opposed to loss-aversive behavior that is seen in the gain condition. Together, these results illustrate that the amount of shift in target selection due to unequal expected reward was suboptimal. As demonstrated below, our modeling results can explain why such optimization is not possible because of the loci of reward influence.

### Saccadic reaction time reflects the effect of task parameters

In our experiments, the subjects were not instructed to saccade as quickly as possible and had to wait until both targets were presented before making a saccade. Nonetheless, we analyzed the saccadic reaction time (SRT) using a stepwise GLM model (see Materials and Methods) to examine whether the SRT reflects any task parameters. The stepwise GLM revealed that the TOA, unequal reward condition, response accuracy, and interaction between the TOA and response accuracy and between the TOA and unequal reward condition had significant effects on the SRT (stepwise GLM: F(5,32394) = 452, *p* = 10^−273^, adjusted-*R*^2^ = 0.065).

First, we found that the SRT decreased with the absolute value of the TOA corresponding to easier trials (*β* for TOA= –1.11, *p* = 0.04; **Fig. 8A**). Second, unequal reward outcomes resulted in an overall decrease in the SRT in the gain and loss conditions compared to the neutral condition (*β* for reward condition= −0.11, *p* = 1.03 × 10^−26^; **Fig. 8A**). Third, the SRT was significantly smaller for correct trials compared to incorrect trials (*β* for response accuracy = −0.076, *p* = 0.0005; **Fig. 8B**). As mentioned above, the stepwise GLM did not reveal a significant effect of the chosen-target relative value on the SRT (**Fig. 8C**). This lack of evidence for a significant effect could be caused by a few factors: 1) the stronger effects of TOA and response accuracy on the SRT, 2) different heuristics used to process gain and loss information, and 3) there was no time pressure in our experiments. Overall, these results show that saccadic reaction time was sensitive to the TOA and unequal expected reward outcomes, indicating that both types of information influenced perceptual choice.

**Figure 8.**
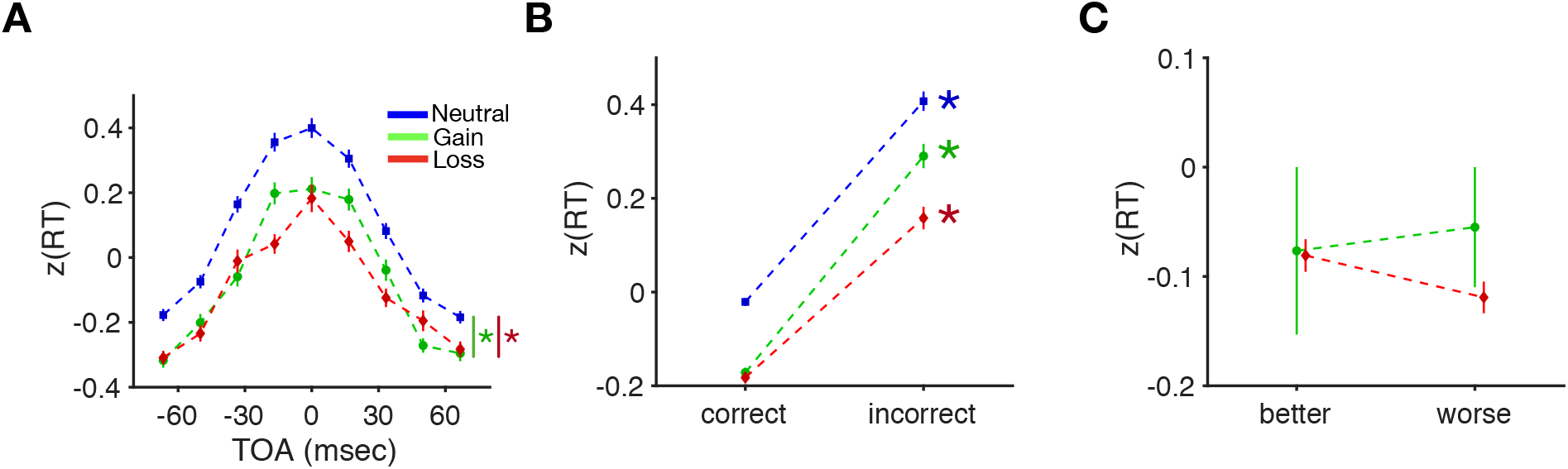
Saccadic reaction time was sensitive to both the TOA and unequal expected reward outcome, reflecting response accuracy, but did not differ between the selection of the better and worse targets. (**A**) Plotted is the z-scored saccadic reaction time (SRT) as a function of the TOA for the neutral (blue), gain (green), and loss (red) conditions. An asterisk shows a significant difference between the average SRT in the neutral and gain, or neutral and loss conditions (stepwise GLM, *p* < 0.05). (**B**) Plotted is the average z-scored SRT on correct and incorrect trials, separately for the three reward conditions indicated in panel A. An asterisk shows a significant difference between the average SRT on correct and incorrect trials (stepwise GLM, *p* < 0.05). (**C**) Plotted is the average SRT on trials in which the chosen target was the target with higher or lower expected reward corresponding to the better and worse targets, respectively.

### Plausible neural mechanisms for observed shifts in target selection

To reveal plausible neural mechanisms underlying the shifts in target selection, we simulated our experimental observations using a cortical network model that we have previously used to successfully simulate the paired-target task (Soltani et al., 2013). Specifically, we focused on capturing our two main experimental findings: 1) similar shifts in target selection during Experiments 1 and 2; 2) lack of correlation between the shifts in target selection and individuals’ sensitivity to sensory evidence in both experiments.

The model consisted of two neural columns with two pools of excitatory neurons and one inhibitory pool of neurons in the superficial layer, and two pools of excitatory neurons in the deep layer (**Supplementary Fig. 1**; see Materials and Methods for more details). To simulate the effect of reward information, we considered three alternative mechanisms that could influence different parts of the model to mimic different stages of decision-making processes: reward-based input to the excitatory pools in the deep layer (**Fig. 9A**; Mechanism 1); reward-dependent gain modulation of sensory input that gives rise to stronger input for the target on the more rewarding side (**Fig. 9B**; Mechanism 2); facilitation of response to the target with higher expected reward in higher visual areas (**Fig. 9B**; Mechanism 3).

**Figure 9.**
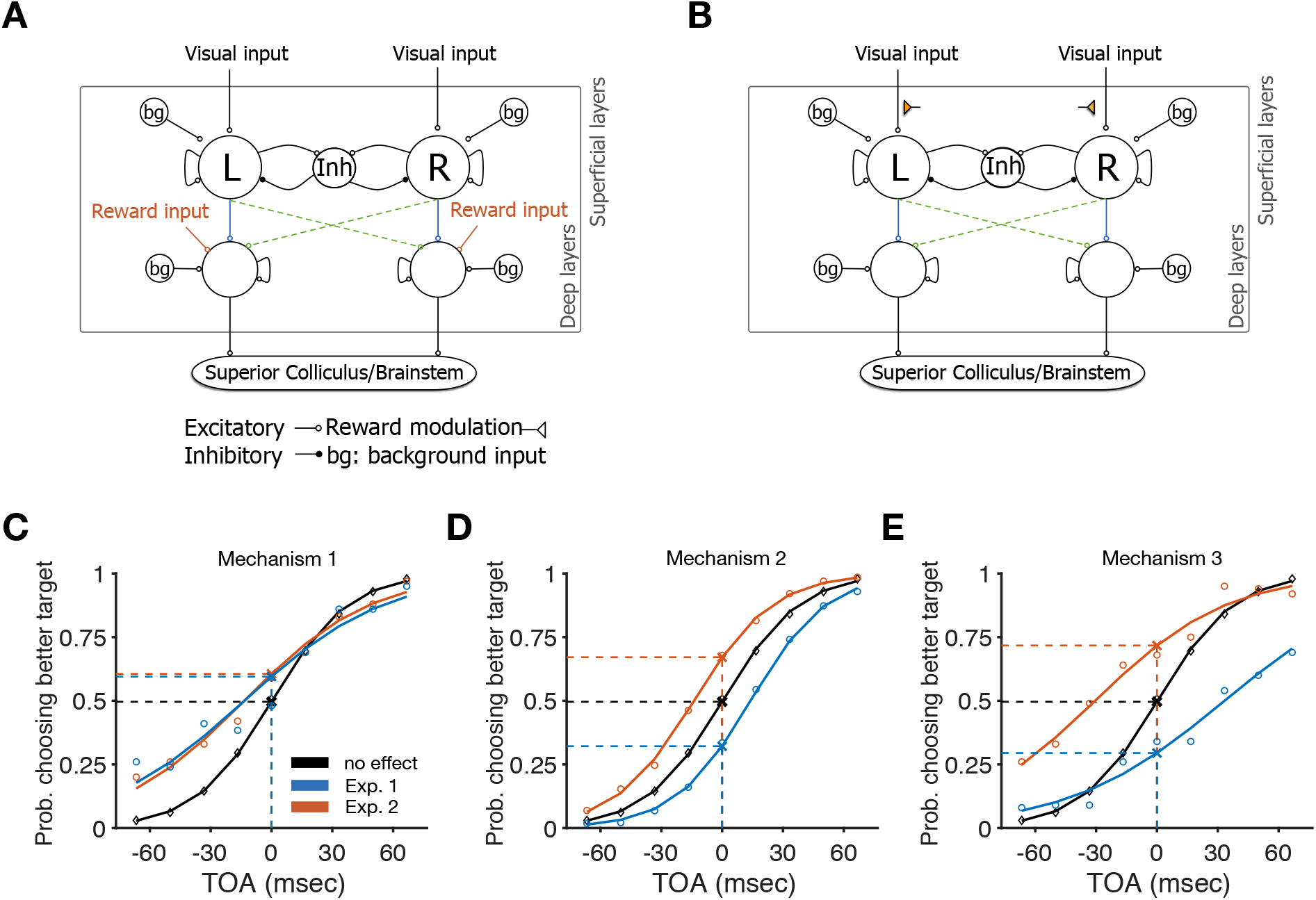
Alternative mechanisms for simulating the effects of unequal expected reward on perceptual decision making. (**A**) The extended model with independent reward input (Mechanism 1). In this model, excitatory pools in deep layer receive an additional reward-based input that was independent of the visual input. The blue solid lines for Experiment 1 and green dashed lines for Experiment 2 show the output projection of excitatory pools of superficial layer to deep layer excitatory pools. (**B**) The extended model with reward-dependent modulation of visual input (Mechanisms 2 and 3). In this model, the visual input to the excitatory pools of superficial layer is modulated by reward information provided at the beginning of the trial. This modulation was performed via two different mechanisms. For Mechanism 2, reward information strengthens (weakens) the visual input for the target with higher (lower) expected reward. In Mechanism 3, unequal reward information results in faster processing (i.e., earlier onset) of the target with higher expected reward. (**C**) Probability of choosing the target on the more rewarding (better) side as a function of TOA in Experiment 1 (and −TOA in Experiment 2) using Mechanism 1. The results with no reward modulation (black diamonds) are shown as the control. The model with Mechanism 1 (circles) produces similar shift in target selection in the two experiments. (**D-E**) Similar to panel C but for the model with Mechanisms 2 (D) and 3 (E). The models with reward-dependent modulation of visual input (Mechanisms 2 and 3) produce opposite shifts in the two experiments.

Simulation results showed that that unequal expected reward results in significant shifts in target selection using all three mechanisms (**Fig. 9C-E**). These shifts, however, were similar for Experiments 1 and 2 only in the model based on Mechanism 1 (**Fig. 9C**). We also examined the correlation between the shift in target selection and the sensitivity to visual input for a given set of model parameters. To generate target selection with different levels of sensitivity to sensory evidence, we changed the background noise in the input to the superficial layer. We did not find any evidence for correlation between the shifts in target selection and sensitivity to the TOA in the model based on Mechanism 1 (Pearson correlation; Experiment 1: *r* = –0.14, *p* = 0.15; Experiment 2: *r* = –0.19, *p* = 0.054; **Fig. 10A, D**). This non-significant correlation, however, had a negative sign similar to that of the experimental data. In contrast, in the model based on Mechanism 2, the shifts in target selection were negatively and positively correlated with sensitivity to the TOA in Experiments 1 and 2, respectively (Pearson correlation; Experiment 1: *r* = –0.58, *p* = 1.75 × 10^−10^; Experiment 2: *r* = 0.56, *p* = 1.46 × 10^−9^; **Fig. 10B, E**). We found qualitatively similar results for the model based on Mechanism 3, even though the shifts in target selection were not significantly correlated to sensitivity to the TOA (Pearson correlation; Experiment 1: *r* = –0.09, *p* = 0.33; Experiment 2: *r* = 0.08, *p* = 0.4; **Fig. 10C, F**).

**Figure 10.**
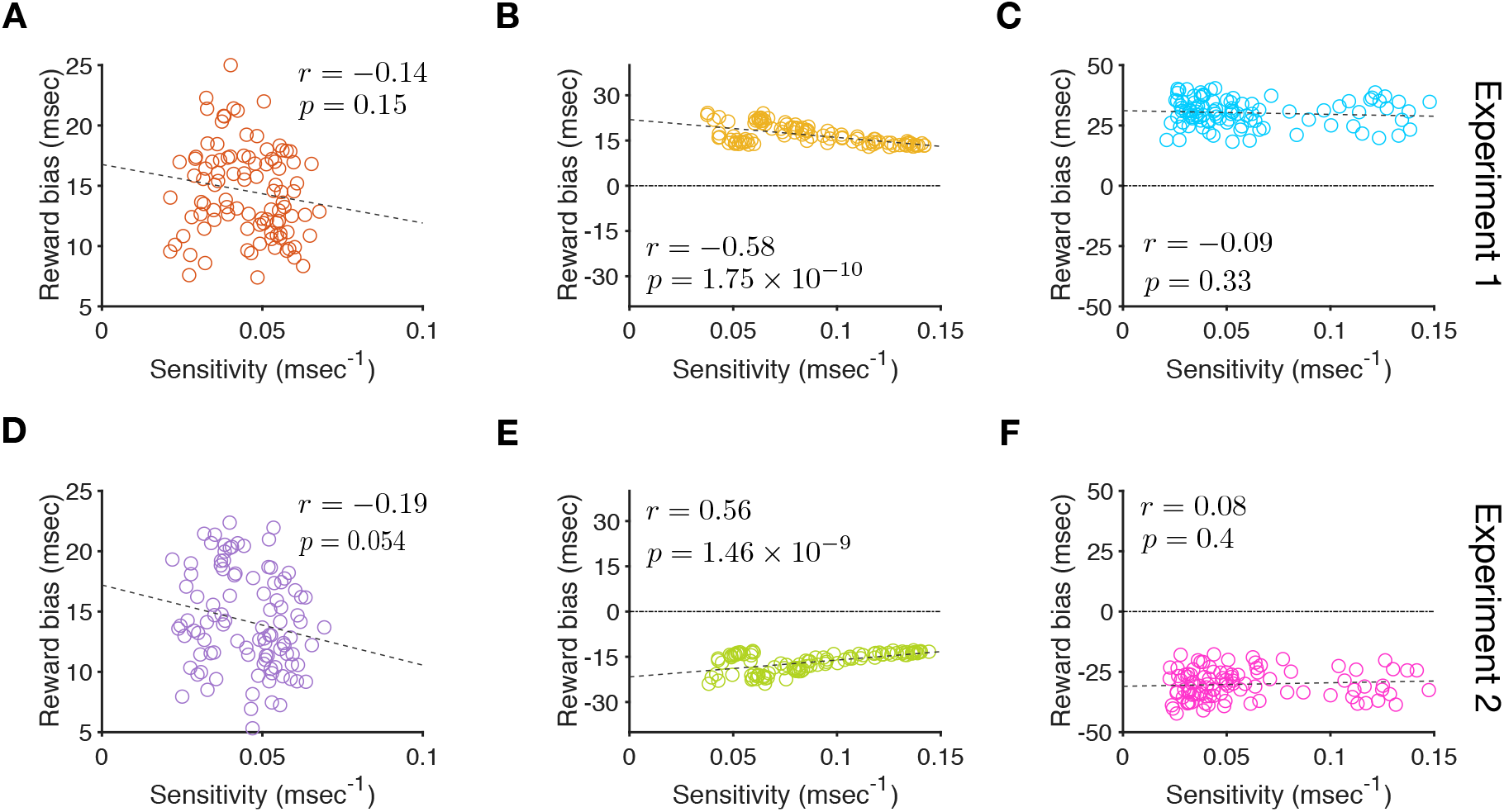
Shift in target selection due to unequal reward input was more compatible with Mechanism 1. (**A**) The shift in target selection is plotted as a function of the sensitivity to the TOA for Experiment 1 using the model based on Mechanism 1. Each point shows the replica of a subject. The grey dashed line shows the least squares line fitted on the simulated behavioral data. (**B–C**) The same as in panel A but using the model based on Mechanisms 2 (B) and 3 (C), respectively. (**D-F**) The same as in panels A-C but for Experiment 2. The models based on Mechanisms 1 and 3 did not show a significant correlation between reward bias and the sensitivity to the TOA. However, the model based on Mechanism 1 did exhibit similar reward bias in the two experiments whereas the model based on Mechanism 3 did not, pointing to Mechanism 1 as the only mechanism that can capture our both main experimental findings.

Overall, these results illustrate that the model based on Mechanism 1 is more compatible with our experimental data for two reasons: 1) exhibiting equal shifts in target selection in Experiments 1 and 2, replicating the observed response bias; and 2) lack of correlation between the shifts in target selection and sensitivity to the TOA in both experiments. These modeling results support the conclusion that the observed shift in subjects’ behavior due to the reward information is more likely to be due to changes in later stages of decision-making. In addition, the modeling results provide a plausible mechanism for how reward information influences perceptual choice. Finally, the results explain why reward biases were independent of sensitivity to sensory signals and thus, could not be optimized.

## Discussion

Series of studies in past decades have aimed to reveal neural mechanisms by which reward influences perceptual decision making. These studies have argued that unequal reward outcomes could either cause a response bias (i.e., increase the tendency to choose the target with larger expected reward) and/or result in a perceptual bias (i.e., differential processing of sensory information). To directly test these two alternative (but not necessarily exclusive) hypotheses for the influence of reward on perceptual decision making, in two sets of experiments, we asked subjects to saccade to the first or second target that appeared on the screen while we manipulated the amount of reward expected from the two alternative responses. Importantly, a bias in sensory processing (i.e., perceptual bias) would result in opposite shifts in target selection in the two experiments, whereas a response bias would cause similar shifts. We did not find any evidence for different amounts of shift in the two experiments, indicating that expected reward is more likely to cause a response bias rather than a bias in sensory processing. These findings dovetail with results from recent studies that used modeling to determine the mechanisms underlying the influence of expected reward on perceptual choice (Diederich, 2008; Diederich & Busemeyer, 2006; Gao et al., 2011) and studies that look at the effect of expectation in general (Bang & Rahnev, 2017; Rungratsameetaweemana, Itthipuripat, Salazar, & Serences, 2018).

Nonetheless, others have argued that reward can directly influence the processing of sensory information during perceptual choice (Cicmil et al., 2015; Liston & Stone, 2008; Pleger et al., 2008; Voss et al., 2008). A possible reason for the discrepancy between their findings and ours could be due to differences in the experimental paradigms in terms of time dependency. The temporal judgment task used here is a type of time-dependent perceptual choice, and it is possible that reward exerts its influence differently during time-independent perceptual choice. For example, the integration of sensory signal over time could push the influence of reward information to later stages of decision making, resulting in response bias instead of perceptual bias. However, there are studies (e.g., Diederich & Busemeyer, 2006) showing the effects of reward as response bias even in discrimination between the lengths of two lines (i.e., time-independent tasks). Either way, future studies are required (using our approach) in order to test the generalizability of our findings to other types of perceptual decision making.

In addition to the reward effects on perception, there is an extensive literature on whether attention influences perception by accelerating sensory processing (the ‘prior entry’ hypothesis) or inducing decision biases. Some of these studies have argued that attention enhances the speed of sensory processing (Hikosaka, Miyauchi, & Shimojo, 1993; Stelmach & Herdman, 1991), whereas other studies have maintained that observed effects are primarily due to attentional modifications of the decision mechanisms (Schneider & Bavelier, 2003). Most of these studies used the so-called temporal order judgment (TOJ) task to measure the point of subjective simultaneity (PSS) from attentional cueing. For example, in a study design similar to ours but concerned with measuring the effects of attention, Shore and colleagues (Shore, Spence, & Klein, 2001) asked subjects to report the first or second targets that appeared on the screen in order to separate changes in sensory processing from response biases. By comparing the shifts in psychometric functions in the two tasks, they found that attention mainly influences perception by accelerating sensory processing in addition to second-order response biases. In contrast, Schneider and Bavelier (Schneider & Bavelier, 2003) have argued that the shift in the PSS due to attentional cueing in the TOJ task is not an adequate reason to accept the prior entry hypothesis. Instead, they suggest that one should compare shifts in the PSS in the TOJ task with those of in a simultaneity judgment task, in which the subjects report whether two stimuli appeared simultaneously or successively. By making this comparison, they showed that attentional cueing has little influence on accelerating sensory processing.

Here, rather than explicit attentional cueing, we used unequal reward information to bias subjects’ processing of information and/or decision making, both of which had behavioral benefits in terms of harvested reward. We also provided reward feedback (correct or incorrect judgment) on each trial, which allowed subjects to correct their biases if desired so. The fact that we observed almost opposite results to those by Shore and colleagues (Shore et al., 2001) based on attentional cueing indicates that reward information influences perception rather differently than how attention affects perception, and therefore, reward and attentional processes rely on different neural mechanisms to guide behavior. Furthermore, because reward information in our experiments was not predictive of the correct response, it is possible that this type of cueing exploits a different mechanism. Nonetheless, attention has been shown to closely interact with reward processing (Farashahi, Azab, Hayden, & Soltani, 2018; Serences, 2008; Soltani, Khorsand, Guo, Farashahi, & Liu, 2016; Spitmaan, Chu, & Soltani, 2019; Stănişor et al., 2013) and revealing that relationship is crucial for fully comprehending both processes (Maunsell, 2004).

It is important to note that even though reward information was not predictive of the correct response in our experiments, subjects still could use this information to obtain more reward. Increasing sensitivity to the more rewarding side does not help detection of the correct target but can improve performance in terms of obtained reward points because temporal judgment is not perfect. For a similar reason, optimal observer such as sequential ratio test can incorporate reward information to adjust decision criterion (Gold & Shadlen, 2001). Although an ideal observer may only change the threshold for response to the more rewarding side, a “less ideal” observer may be persuaded to attend more to the more rewarding side, which could result in a change in perception.

To reveal possible neural mechanisms underlying our observations, we extended our biophysically-plausible cortical network model (Soltani et al., 2013) to simulate shifts in target selection due to unequal expected reward based on alternative mechanisms. We found that our experimental results are more compatible with the influence of reward information on later stages of decision-making processes via biasing the activity in the output layer of the decision circuit toward the target on the more rewarding side. Considering that lesions and reversible inactivation of the frontal eye field (FEF) cause similar shifts in target selection during the paired-target task (Schiller & Chou, 1998, 2000; Schiller & Tehovnik, 2003), reward could exerts its influence through modulations of the output layer of the FEF. Furthermore, our analyses of saccadic reaction time revealed that unequal expected reward outcomes resulted in faster decision making. Future experiments that emphasize speed could provide additional information to test alternative models.

Importantly, we found that to reproduce our experimental results by our model, the input for biasing target selection should be independent of sensory evidence, which is more consistent with results from a few recent studies (Diederich, 2008; Diederich & Busemeyer, 2006; Gao et al., 2011). For example, using a task in which the subjects had to judge whether two lines are the same or different while manipulating the payoff for the two responses, Diederich and Busemeyer have shown that the effect of unequal payoffs is more compatible with a two-stage processing of sensory and reward information (Diederich & Busemeyer, 2006). In their model, the decision maker first integrates reward information followed by the integration of sensory information (with no reward modulation) if no decision is made during the first stage. In another study, Gao and colleagues used a leaky competing-accumulator model to show that reward information biases the initial state of the decision variable toward the target with higher expected reward (Gao et al., 2011). Both these studies illustrate that reward information does not interact with sensory evidence. In contrast, in our study, all of the mechanisms explored generated comparable shifts in target selection in Experiment 1 suggests that in order to distinguish the origin of reward effects, one needs to consider the appropriate task design in addition to the appropriate model.

Our results not only show that the amounts of shifts in target selection due to unequal expected reward were suboptimal and independent of individuals’ sensitivity to sensory signal, but also explain that these shifts could not be optimized due to reward influence on later stages of decision making. In addition, we observed a larger shift in target selection in the loss condition compared to gain condition in both experiments. This result resembles loss-aversion behavior during value-based choice (Tversky & Kahneman, 1992) and extends this phenomenon to perceptual decision making with different reward outcomes. Together, these findings suggest that even during perceptual choice, there are some heuristics in differential processing of gain and loss information.

Similar but much weaker suboptimal behavior has also been observed for biased reward probabilities during perceptual decision making (Navalpakkam, Koch, & Perona, 2009; Voss et al., 2008). Interestingly, humans exhibit a closer-to-optimal criterion when they deal with unequal reward probabilities rather than unequal reward magnitudes on alternative options or actions (Maddox, 2002; Teichert & Ferrera, 2010). Our modeling results indicate that shifts in target selection can be closer to optimal if reward information affects the processing of visual input instead of the final stages of decision making. Therefore, the difference in response to unequal reward probability and magnitude could be due to their influence on different stages of decision making. Finally, in environments that resemble more naturalistic settings, adjustments in choice and learning of reward probability can occur in the absence of any optimization (Farashahi et al., 2017; Khorsand & Soltani, 2017). Future studies are required to determine whether reward probability and magnitude exert their influence at separate stages of decision making.

## Acknowledgments

We would like to thank Daeyeol Lee, Shiva Farashahi, and Mehran Spitmaan for help in early stages of this work, and Patrick Cavanagh and Shih-Wei Wu for comments on the manuscript. This work was supported by National Science Foundation (EPSCoR Award 1632738 to A.S.).

## Supplementary figures

**Supplementary Figure 1.**
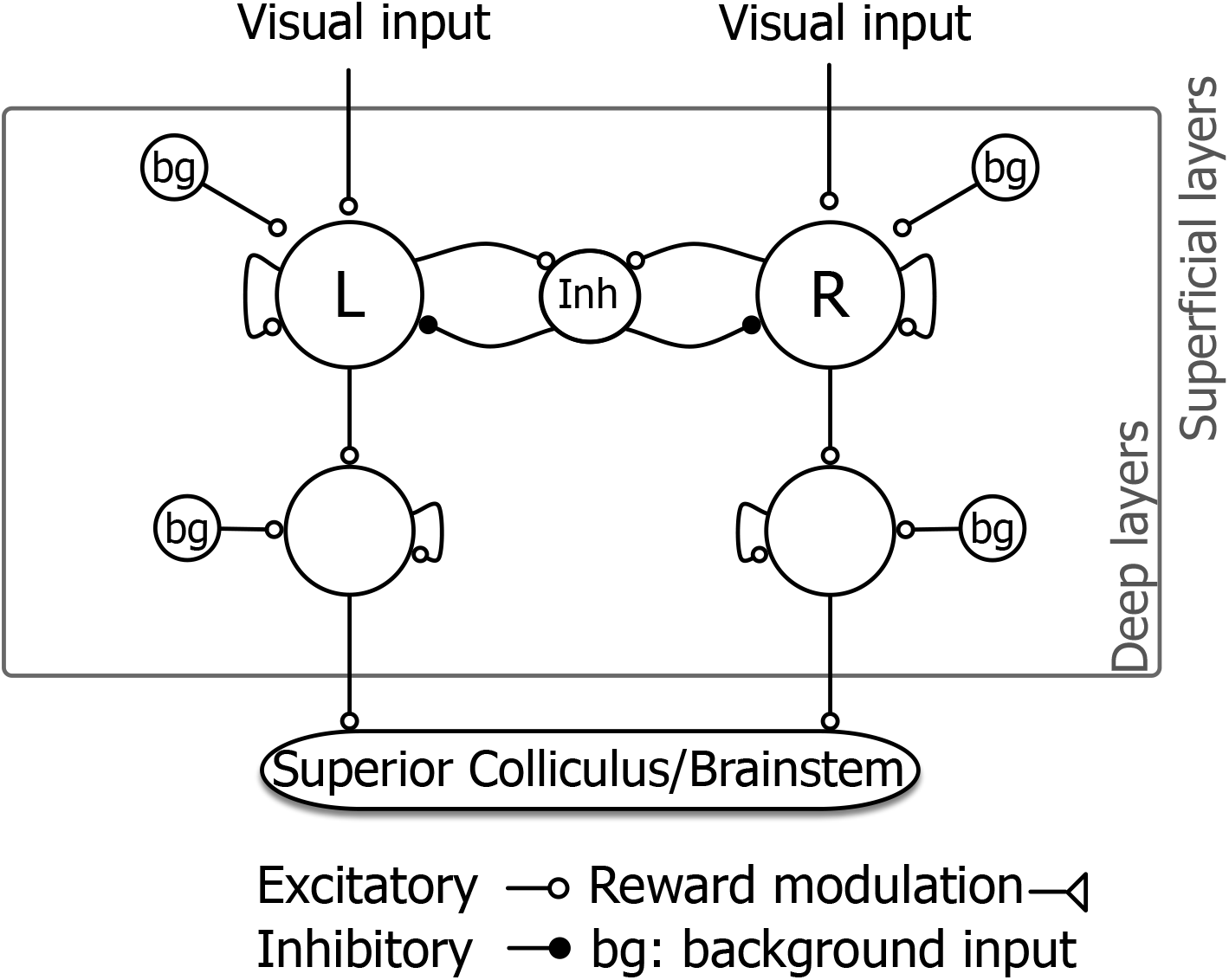
The basic architecture of the original network model used to simulate the paired-target task.

**Supplementary Figure 2.**
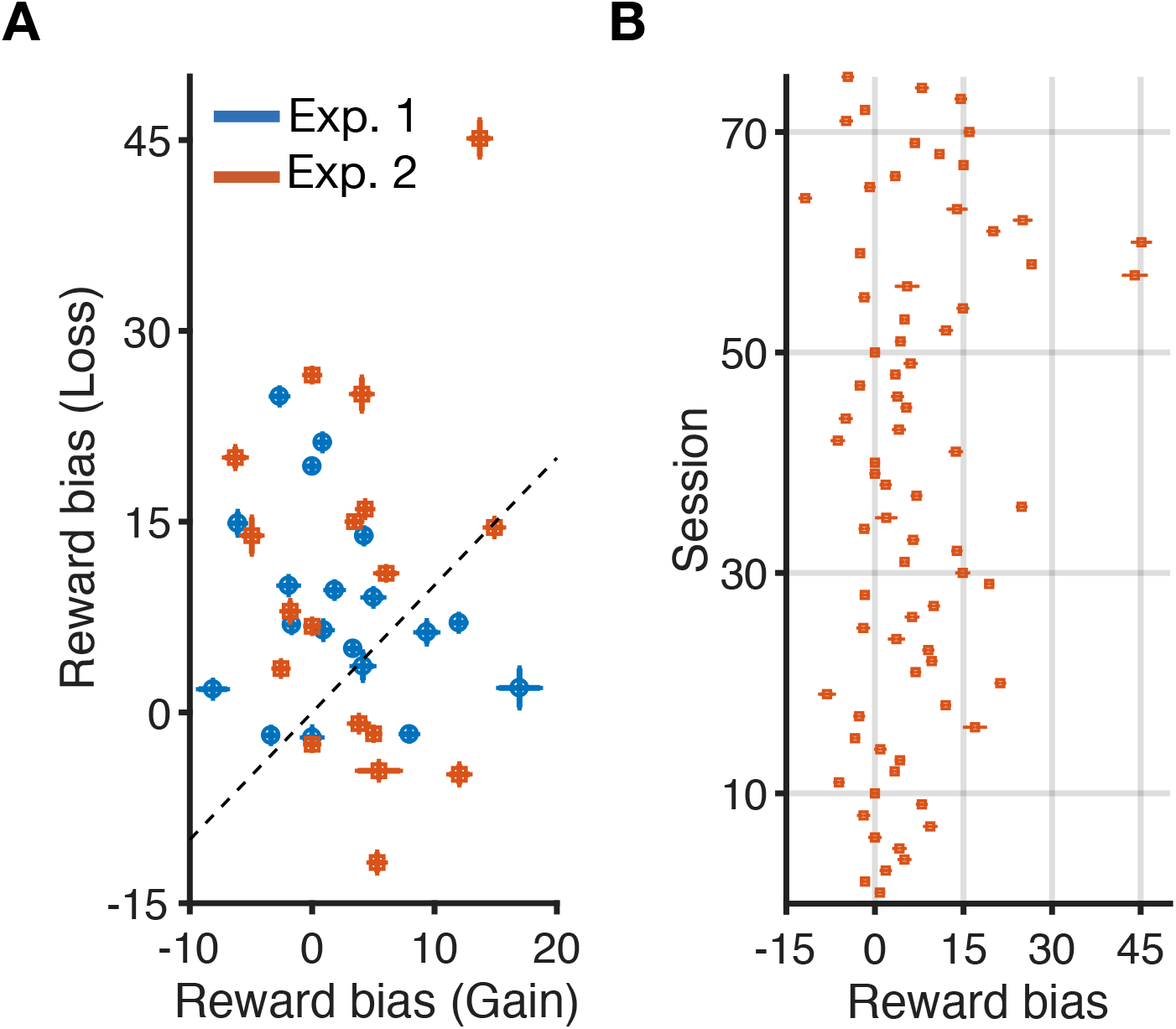
Robustness of the estimated reward biases. (**A**) Plotted are reward biases in the loss condition as a function of the reward biases in the gain condition, separately for Experiments 1 (blue) and 2 (red). Error bars show 95% confidence intervals and are calculated based on the Hessian matrix of the log-likelihood function. In most cases, the error bars are smaller than the symbol. (**B**) Plotted are reward biases and their estimation error for all sessions of the two experiments (in both loss and gain conditions).

**Supplementary Figure 3.**
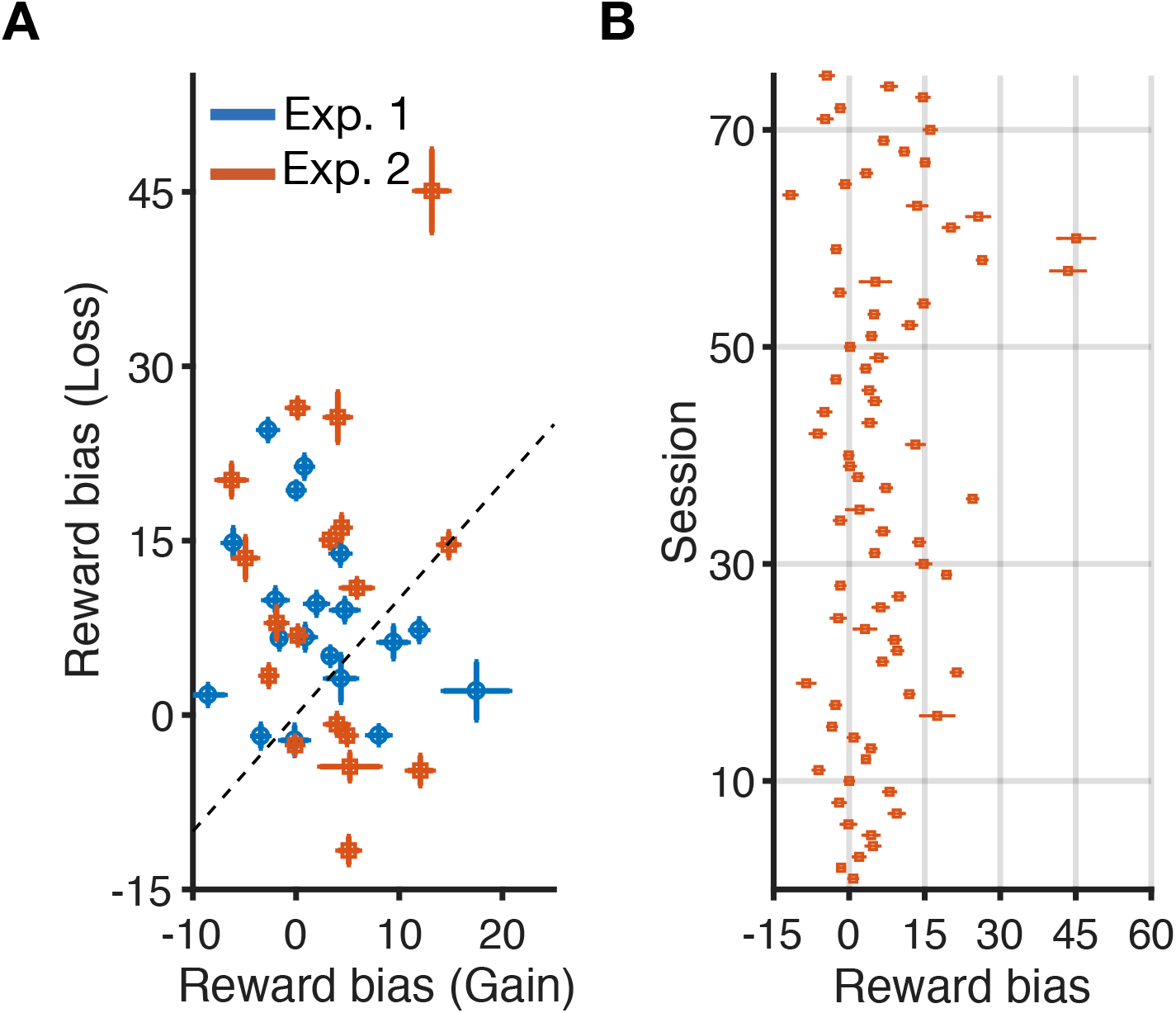
Robustness of the estimated reward biases. (**A**) Plotted are reward biases in the loss condition as a function of the reward biases in the gain condition, separately for Experiments 1 and 2. Error bars are standard deviation calculated based on sampling 95% of the data 50 times and fitting the ensuing psychometric function. (**B**) Plotted are reward biases and their estimation error for all sessions of the two experiments (in both loss and gain conditions).

## References

Bang, J. W., & Rahnev, D. (2017). Stimulus expectation alters decision criterion but not sensory signal in perceptual decision making. Scientific Reports, 7(1), 1–12. https://doi.org/10.1038/s41598-017-16885-2

Brainard, D. H. (1997). The Psychophysics toolbox. Spatial Vision, 10(4), 433–436. https://doi.org/10.1163/156856897X00357

Carrasco, M., & Barbot, A. (2019). Spatial attention alters visual appearance. Current Opinion in Psychology, 29, 56–64. Retrieved from https://doi.org/10.1016/j.copsyc.2018.10.010

Carrasco, M., Ling, S., & Read, S. (2004). Attention alters appearance. Nature Neuroscience, 7(3), 308–313. https://doi.org/10.1038/nn1194

Christopoulos, V., Bonaiuto, J., & Andersen, R. A. (2015). A biologically plausible computational theory for value integration and action selection in decisions with competing alternatives. PLoS Computational Biology, 11(3), 1–31. https://doi.org/10.1371/journal.pcbi.1004104

Christopoulos, V., & Schrater, P. R. (2015). Dynamic integration of value information into a common probability currency as a theory for flexible decision making. PLoS Computational Biology, 11(9), 1–26. https://doi.org/10.1371/journal.pcbi.1004402

Cicmil, N., Cumming, B. G., Parker, A. J., & Krug, K. (2015). Reward modulates the effect of visual cortical microstimulation on perceptual decisions. ELife, 4(September), 1–25. https://doi.org/10.7554/eLife.07832

Diederich, A. (2008). A further test of sequential-sampling models that account for payoff effects on response bias in perceptual decision tasks. Perception & Psychophysics, 70(2), 229–256. https://doi.org/10.3758/PP.70.2.229

Diederich, A., & Busemeyer, J. R. (2006). Modeling the effects of payoff on response bias in a perceptual discrimination task: Bound-change, drift-rate-change, or two-stage-processing hypothesis. Perception and Psychophysics, 68(2), 194–207. https://doi.org/10.3758/BF03193669

Farashahi, S., Azab, H., Hayden, B., & Soltani, A. (2018). On the flexibility of basic risk attitudes in monkeys. The Journal of Neuroscience, 38(18), 4383–4398. https://doi.org/10.1523/JNEUROSCI.2260-17.2018

Farashahi, S., Donahue, C. H., Khorsand, P., Seo, H., Lee, D., & Soltani, A. (2017). Metaplasticity as a neural substrate for adaptive learning and choice under uncertainty. Neuron, 94(2), 401–414.e6. https://doi.org/10.1016/j.neuron.2017.03.044

Farashahi, S., Ting, C. C., Kao, C. H., Wu, S. W., & Soltani, A. (2018). Dynamic combination of sensory and reward information under time pressure. PLoS Computational Biology, 14(3), 1–26. https://doi.org/10.1371/journal.pcbi.1006070

Feng, S., Holmes, P., Rorie, A., & Newsome, W. T. (2009). Can monkeys choose optimally when faced with noisy stimuli and unequal rewards? PLoS Computational Biology, 5(2). https://doi.org/10.1371/journal.pcbi.1000284

Gao, J., Tortell, R., & McClelland, J. L. (2011). Dynamic integration of reward and stimulus information in perceptual decision-making. PLoS ONE, 6(3), 5–7. https://doi.org/10.1371/journal.pone.0016749

Gold, J., & Shadlen, M. (2001). Neural computations that underlie decisions about sensory stimuli. TRENDS in Cognitive Sciences, 5(1), 10–16.

Hikosaka, O., Miyauchi, S., & Shimojo, S. (1993). Focal visual attention produces illusory temporal order and motion sensation. Vision Research, 33(9), 1219–1240. https://doi.org/10.1016/0042-6989(93)90210-N

Khorsand, P., & Soltani, A. (2017). Optimal structure of metaplasticity for adaptive learning. PLoS Computational Biology, 13(6), 1–22. https://doi.org/10.1371/journal.pcbi.1005630

Kleiner, M., Brainard, D. H., Pelli, D. G., Broussard, C., Wolf, T., & Niehorster, D. (2007). What’s new in Psychtoolbox-3? Perception, 36, S14. https://doi.org/10.1068/v070821

Liston, D. B., & Stone, L. S. (2008). Effects of prior information and reward on oculomotor and perceptual choices. Journal of Neuroscience, 28(51), 13866–13875. https://doi.org/10.1523/JNEUROSCI.3120-08.2008

Maddox, W. T. (2002). Toward a unified theory of decision criterion learning in perceptual categorization. Journal of the Experimental Analysis of Behavior, 78(3), 567–595. https://doi.org/10.1901/jeab.2002.78-567

Maunsell, J. H. R. (2004). Neuronal representations of cognitive state: reward or attention? Trends in Cognitive Sciences, 8(6), 261–265. https://doi.org/10.1016/j.tics.2004.04.003

Mulder, M. J., Wagenmakers, E.-J., Ratcliff, R., Boekel, W., & Forstmann, B. U. (2012). Bias in the brain: a diffusion model analysis of prior probability and potential payoff. Journal of Neuroscience, 32(7), 2335–2343. https://doi.org/10.1523/JNEUROSCI.4156-11.2012

Navalpakkam, V., Koch, C., & Perona, P. (2009). Homo economicus in visual search. Journal of Vision, 9(1)(31), 1–16. https://doi.org/10.1167/9.1.31.Introduction

Pelli, D. G. (1997). The VideoToolbox software for visual psychophysics: Transforming numbers into movies. Spatial Vision. https://doi.org/10.1163/156856897X00366

Pleger, B., Blankenburg, F., Ruff, C. C., Driver, J., & Dolan, R. J. (2008). Reward facilitates tactile judgments and modulates hemodynamic responses in human primary somatosensory cortex. Journal of Neuroscience, 28(33), 8161–8168. https://doi.org/10.1523/JNEUROSCI.1093-08.2008

Rajsic, J., Perera, H., & Pratt, J. (2017). Learned value and object perception: Accelerated perception or biased decisions? Attention, Perception, and Psychophysics, 79(2), 603–613. https://doi.org/10.3758/s13414-016-1242-0

Rorie, A. E., Gao, J., McClelland, J. L., & Newsome, W. T. (2010). Integration of sensory and reward information during perceptual decision-making in Lateral Intraparietal Cortex (LIP) of the macaque monkey. PLoS ONE, 5(2). https://doi.org/10.1371/journal.pone.0009308

Rungratsameetaweemana, N., Itthipuripat, S., Salazar, A., & Serences, J. T. (2018). Expectations do not alter early sensory processing during perceptual decision making. The Journal of Neuroscience, 3638–17. https://doi.org/10.1523/JNEUROSCI.3638-17.2018

Schiller, P. H., & Chou, I. H. (1998). The effects of frontal eye field and dorsomedial frontal cortex lesions on visually guided eye movements. Nat Neurosci, 1(3) 248–253. https://doi.org/10.1038/693

Schiller, P. H., & Chou, I. H. (2000). The effects of anterior arcuate and dorsomedial frontal cortex lesions on visually guided eye movements: 2. Paired and multiple targets. Vision Research, 40(10–12), 1627–1638. https://doi.org/10.1016/S0042-6989(00)00058-4

Schiller, P. H., & Tehovnik, E. J. (2003). Cortical inhibitory circuits in eye-movement generation. European Journal of Neuroscience, 18(11), 3127–3133. https://doi.org/10.1111/j.1460-9568.2003.03036.x

Schneider, K. A., & Bavelier, D. (2003). Components of visual prior entry. Cognitive Psychology, 47(4), 333–366. https://doi.org/10.1016/S0010-0285(03)00035-5

Serences, J. T. (2008). Value-based modulations in human visual cortex. Neuron, 60(6), 1169–1181. https://doi.org/10.1016/j.neuron.2008.10.051

Shore, D. I., Spence, C., & Klein, R. M. (2001). Visual prior entry. Psychological Science, 12(3), 205–212. Retrieved from http://www.ijrap.net/admin/php/uploads/1308_pdf.pdf

Soltani, A., Khorsand, P., Guo, C., Farashahi, S., & Liu, J. (2016). Neural substrates of cognitive biases during probabilistic inference. Nature Communications, 7, 1–14. https://doi.org/10.1038/ncomms11393

Soltani, A., Noudoost, B., & Moore, T. (2013). Dissociable dopaminergic control of saccadic target selection and its implications for reward modulation. Proceedings of the National Academy of Sciences, 110(9), 3579–3584. https://doi.org/10.1073/pnas.1221236110

Spitmaan, M., Chu, E., & Soltani, A. (2019). Salience-driven value construction for adaptive choice under risk. The Journal of Neuroscience, 39(26), 2522–18. https://doi.org/10.1523/jneurosci.2522-18.2019

Stanford, T. R., Shankar, S., Massoglia, D. P., Costello, M. G., & Salinas, E. (2010). Perceptual decision making in less than 30 milliseconds. Nature Neuroscience, 13(3), 379–385. https://doi.org/10.1038/nn.2485

Stănişor, L., van der Togt, C., Pennartz, C. M. A., Roelfsema, P. R., Stanisor, L., van der Togt, C., … Roelfsema, P. R. (2013). A unified selection signal for attention and reward in primary visual cortex. Proceedings of the National Academy of Sciences, 110(22), 9136–9141. https://doi.org/10.1073/pnas.1300117110

Stelmach, L. B., & Herdman, C. M. (1991). Directed attention and perception of temporal order. Journal of Experimental Psychology: Human Perception and Performance, 17(2), 539–550. https://doi.org/10.1037/0096-1523.17.2.539

Sugrue, L. P., Corrado, G. S., & Newsome, W. T. (2005). Choosing the greater of two goods: Neural currencies for valuation and decision making. Nature Reviews Neuroscience, 6(5), 363–375. https://doi.org/10.1038/nrn1666

Summerfield, C., & Koechlin, E. (2010). Economic value biases uncertain perceptual choices in the parietal and prefrontal portices. Frontiers in Human Neuroscience, 4(November), 1–12. https://doi.org/10.3389/fnhum.2010.00208

Teichert, T., & Ferrera, V. P. (2010). Suboptimal integration of reward magnitude and prior reward likelihood in categorical decisions by monkeys. Frontiers in Neuroscience, 4(NOV), 1–13. https://doi.org/10.3389/fnins.2010.00186

Tosoni, A., Committeri, G., Calluso, C., & Galati, G. (2017). The effect of reward expectation on the time course of perceptual decisions. European Journal of Neuroscience, 45(9), 1152–1164. https://doi.org/10.1111/ejn.13555

Tversky, A., & Kahneman, D. (1992). Advances in prospect theory: cumulative representation of uncertainty. Journal of Risk and Uncertainty, 5, 297–323. https://doi.org/10.1016/j.actao.2004.05.005

Voss, A., Rothermund, K., & Brandtstädter, J. (2008). Interpreting ambiguous stimuli: Separating perceptual and judgmental biases. Journal of Experimental Social Psychology, 44(4), 1048–1056. https://doi.org/10.1016/j.jesp.2007.10.009

Wong, K.-F., & Wang, X.-J. (2006). A recurrent network mechanism of time integration in perceptual decisions. Journal of Neuroscience, 26(4), 1314–1328. https://doi.org/10.1523/JNEUROSCI.3733-05.2006

